# Telomere-to-telomere genome of the model plant *Physcomitrium patens*

**DOI:** 10.1101/2023.05.19.541548

**Authors:** Guiqi Bi, Shijun Zhao, Jiawei Yao, Huan Wang, Mengkai Zhao, Yuanyuan Sun, Xueren Hou, Yuling Jiao, Yingxin Ma, Jianbin Yan, Junbiao Dai

## Abstract

The model plant *Physcomitrium patens* (*P. patens*) has played a pivotal role in enhancing our comprehension of plant evolution, growth, and development. However, the current genome harbors numerous intricate regions that remain unfinished and erroneous. To address these issues, we present an exemplary assembly of the *P. patens* nuclear genome, which incorporates telomeres and centromere regions, thereby establishing it as the telomere-to-telomere (T2T) genome in a non-seed plant. This T2T genome not only dispels the prevailing misconception regarding chromosome number in *P. patens* but also provides indispensable resources for conducting in-depth studies in moss genomics and biology.

## Main

The moss *Physcomitrium* (*Physcomitrella*) *patens* has been utilized as an experimental plant for more than half a century due to its exceptional properties, which include low morphological complexity, a high frequency of homologous recombination, and high tolerance to stress. Furthermore, it offers established tools and resources for genetic analysis and manipulation^1–4^, rendering it a model organism for investigating various aspects of plant biology, such as plant terrestrialization, non-seed plant evolution, tip cell growth, stem cell formation, and cell metabolism^5–9^.

*P. patens* occupies a significant phylogenetic position as a result of its classification as a member of the bryophytes, the sister lineage to vascular plants. This led to its selection as the primary non-seed plant for genome sequencing. The draft of its nuclear genome was initially obtained through Sanger sequencing in 2008^10^, enabling it to be included in genome-era research alongside model plants such as *Arabidopsis*^11^, rice^12^, and *Chlamydomonas*^13^. By utilizing a genetic linkage map and implementing Sanger sequencing techniques, a substantial improvement was made to the genome assembly, leading to its promotion to the chromosomal level (version V3). This updated version exhibits an impressive contig N50 of 465 kbp and has effectively anchored 462.3 Mbp of genome sequences onto 27 chromosomes^14^.

The inclusion of annotations in the moss genome has facilitated and expedited the investigation of genomic and molecular aspects of *P. patens*. While the reference genome provides a broad overview of chromosome structures, it is still riddled with inaccuracies and incomplete segments, with 3,940 gaps present throughout the genome. With the development of genome sequencing technologies, achieving telomere-to-telomere (T2T) genome assembly of model organisms has become an essential goal for improving the underlying data source in biological research. Excitingly, the complete human genome^15^, two gap-free rice reference genomes^16–17^, and genomes of kiwi fruit^18, 19^, watermelon^20^ and *Chlamydomonas reinhardtii*^21^ have just been reported successively. With momentous guidance and significance to moss biology, deciphering the complete genome of *P. patens* will provide critical information to meet the requirements of in-depth moss studies.

### Implementation of T2T Assembly

To complete the genome of *P. patens*, we undertook sequencing of the haplotype of the Gransden strain. This led to the acquisition of 5,233,416 Oxford Nanopore MinION reads that encompassed 42.98 Gbp of sequence, with an average length of 8,213 bp. Additionally, we obtained 307,136,852 Illumina NovaSeq read pairs, which amounted to 51.14 Gbp, and 32.08 Gbp of Hi-C data for scaffolding (Supplementary Table 1). Our selection of the genome backbone involved a meticulous evaluation of the assembly results obtained from NextDenovo^22^ and Flye^23^, which served as our primary assembly software. We utilized six different data combinations, which entailed filtering out organelle reads and utilizing various correction software (Supplementary Table 2). After a thorough comparison of the assembly results, we ultimately opted for the NextDenovo assembly results that utilized all ONT reads as the initial backbone. This particular assembly exhibited an N50 of 15.43 Mbp and comprised 45 contigs. Next, we conducted Hi-C scaffolding and manually corrected misassemblies, leading to the development of 26 distinct chromosome structures. Among these, 10 were devoid of gaps, while the other chromosomes had 23 gaps. We also detected a total of 47 telomere locations. Notably, this genome exhibited an unprecedented telomere composition, (GGGCCA)_n_, in addition to the conventional (TTTAGGG)_n_ telomere sequence (Fig. 1a). Given that this novel telomere sequence of *P. patens* is not documented in the telomere database^24^, it can be inferred that the telomere structure of this species is unique and distinct.

**Fig. 1.**
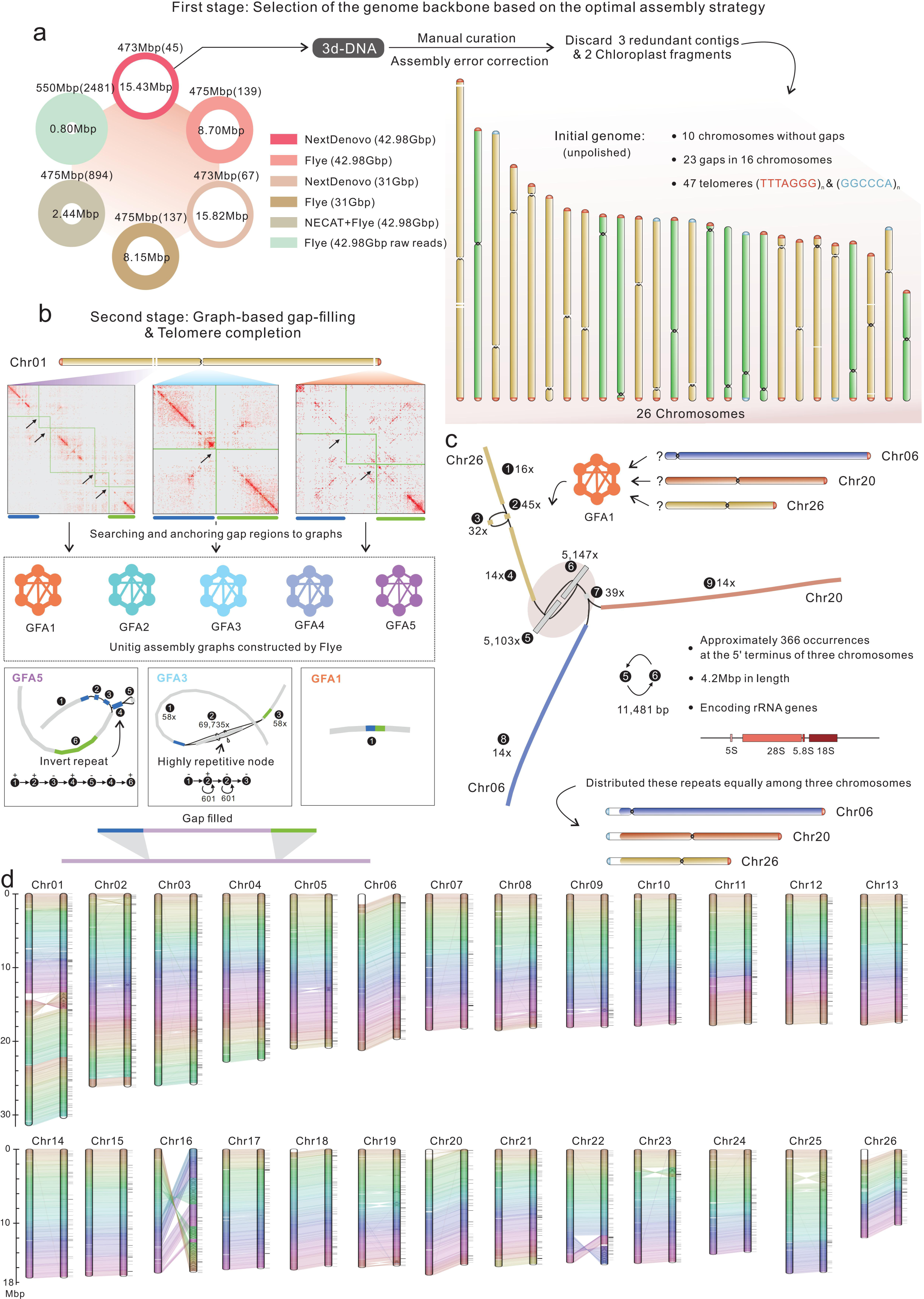
Telomere-to-Telomere (T2T) assembly strategy for the *P. patens* V4 genome. **a**, Selection of the genome backbone based on comparison of assemblies obtained from various combinations of assemblers and datasets and construction of the initial 26 chromosomes through Hi-C scaffolding. Within the display, six assemblies are presented in varying circles situated in the upper left corner. The size of the genomic assembly and N50 values are depicted by the outer and inner circle dimensions, respectively, which are proportional to their corresponding values. The numerical values contained in brackets indicate the numbers of contigs obtained in the corresponding assembly. The NextDenovo assembly, using whole nanopore reads, resulted in 45 contigs. This assembly was chosen for chromosome scaffolding, followed by manual curation. After this procedure, five redundant fragments were removed, leading to 26 chromosomes containing 23 gaps and two types of telomere sequences. **b**, Implementation of the graph-based gap-filling method. We initially generated five graphical fragment assembly (GFA) files utilizing Flye software. These files were created from diverse datasets to address potential inconsistencies in assembly that may have arisen from varying read correction methods. Subsequently, we utilized chromosome 1 as an example, which encompasses three gap intervals with a total of seven gaps. A sequence spanning approximately 200 kbp upstream (blue bar) and downstream (light green bar) of each gap was extracted and mapped to these GFA graphs to find possible reasons causing breaks. Then, an appropriate connection pathway was inferred to fill each gap. **c**, The three-body problem: highly tandem repeats located at the termini of three chromosomes encoding missing rRNA genes. The GFA graph depicts a scenario where three chromosomes are entangled due to the presence of two highly repetitive fragments. Based on the depth information analysis, it has been inferred that the recurrence of these reiterated fragments is 366 times greater than that of typical chromosomes, indicating a comprehensive sequence length of up to 4.2 Mbp. This length exceeds the current capacity of third-generation sequencing techniques. Therefore, a prudent approach would be to distribute this highly repetitive fragment evenly among the three chromosomes. **d**, Genome synteny comparison between the final finished V4 and V3 assemblies. The paired homologous chromosomes are represented by the left bar indicating the V4 genome and the right bar indicating the V3 genome. The V3 genome is distinguished by slim black lines adjacent to it, indicating the position of gaps. To maintain clarity, Chr25 and Chr27 in the V3 version have been merged together, as Chr27 is actually a component of Chr25 in the V4 version.

We have devised the Graph-Based Gap Filling (GBGF) approach to supplement the T2T assembly. This method involves a semimanual process of filling gaps in the Unitig graph by identifying breakpoints and analyzing their underlying causes. Unlike overlap-based filling methods, GBGF meticulously follows the structural characteristics of breakpoints, such as the presence of reverse and tandem repeat structures or complex regions. By constructing precise sequence paths, the gap region is replaced to complete filling, rather than simply filling it mechanically. GBGF has displayed impressive results in testing existing diploid heterozygous genomes (unpublished). Prior to implementing the GBGF strategy, to mitigate discrepancies in outcomes resulting from various read correction algorithms, we concurrently constructed five Unitig graphs (GFA1-5) during the assembly of *P. patens* (Supplementary Table 3). As an example, Chr01 exhibits seven gaps in three broken regions. To scrutinize the formation of these broken sites, we extracted the sequences situated upstream and downstream (50-200 kbp) of these regions and anchored them onto these graphs. Based on the search, we determined that the initial area showcases a reverse-repeating fragment, while the subsequent area showcases highly tandem repeats, and the final area presents a fracture that has arisen due to sequence redundancy. By constructing the path (based on the mapping of long ONT reads and estimating the number of copies), we obtained the completed region sequences, and gap filling was accomplished through sequence replacement (Fig. 1b). For the 16 gaps that remain, 14 of them display constructions adjacent to the centromeres, yet they can be completed with GBGF. The presence of two unfilled gaps on chromosomes 21 and 24 indicates the existence of strikingly similar regions. This, in turn, implies an intertwining of the graphs of the two chromosomes. After deducing the correct path to connect these breakpoints through the mapping of long reads, we achieved the final gap-free assembly (Supplementary Fig. 3).

Two methods were employed to complete the structure of telomeres. The initial approach entailed utilizing GBGF to generate a graph for the telomere site and ascertain the indispensability of the telomere sequence for completion. The subsequent method involved searching for corrected reads to finalize the repair. During this process, it was discovered that the 5’ ends of Chr06, Chr20, and Chr26 were linked due to two highly repetitive fragments, creating a “three-body problem” in the assembly process (Fig. 1c). These two fragments, with a total length of 11,481 bp, occurred approximately 366 times more frequently than other normal fragments in the nuclear genome. This indicates that the total length of these high-copy fragments in the genome is approximately 4.2 Mbp, which surpasses the current long read sequencing capability. Furthermore, this pair of fragments is in the coding region of the rRNA gene, suggesting that the number of rRNA genes encoded was significantly underestimated in previous genome versions. It is plausible that the high tandem repeats of rRNA were uniformly dispersed across the three chromosomes, and the completion of the three telomeres, with (GGGCCA)_n_, was accomplished through the search for corrected ONT reads. The latest telomere-to-telomere assembly, named the V4 version, has a length of 481,750,213 bp and contig N50 of 17.71 Mbp and was obtained after polishing with third- and second-generation sequencing reads (Fig. 1d, Fig. 2a, Supplementary Table 4).

**Fig. 2.**
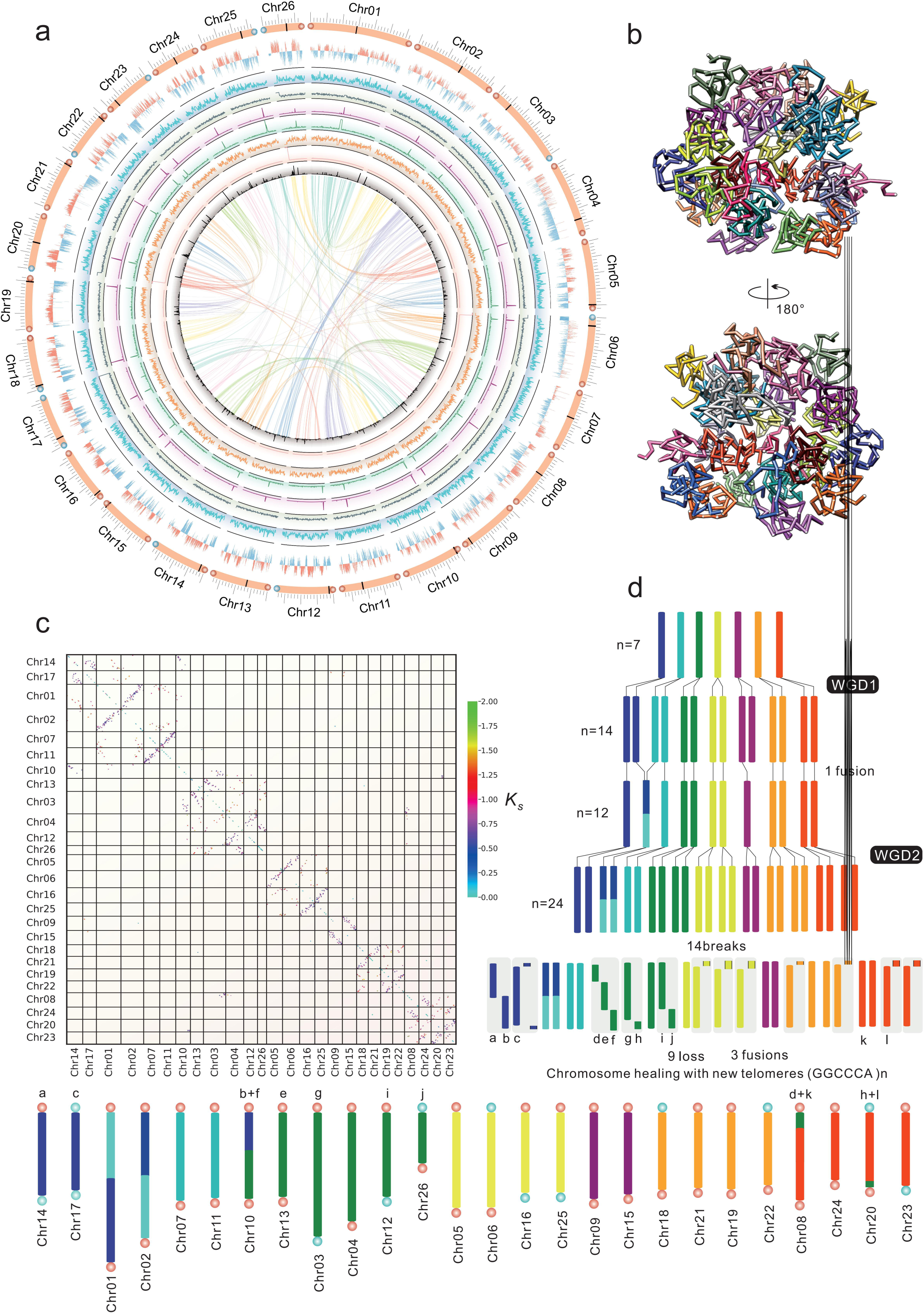
Chromosomal features of the V4 genome. **a**, Genomic landscape of the twenty-six T2T chromosomes. The outermost track exhibits the chromosome karyotype, delineating the centromere positions with black bars. The telomere configuration is represented by dots at both ends of each chromosome, with the conventional telomere sequence (TTTAGGG)_n_ and the novel telomere sequence (GGCCCA)_n_ indicated by red and blue colors, respectively. The second inner track displays the fraction of the genome present in A (pink) or B (light blue) compartments. The subsequent tracks, with a resolution of 100 kbp, depict the density of genes, GC content, centromere-specific Copia density, overall Copia distribution, Gypsy distribution level, rRNA gene density, and tRNA density. The central region emphasizes the collinear regions detected within the genome. **b**, A whole-genome three-dimensional model showing individually colored chromosomes with nodes defining topologically associating domains (TADs). **c**, Dot plot showing two WGD events in *P. patens*. The coloring of dots representing the positions of paralogous gene pairs was determined by the *K_s_* scale. The chromosomes are ordered based on the derivation of 7 ancestral chromosomes. **d**, Updated karyotype evolutionary scenario of *P. patens* based on 26 chromosomes from the ancestral karyotype, illustrated with seven colors. The bottommost segment of the diagram exhibits the present-day genome, which is distinguished by a spectrum of colors that indicate its provenance from the ancestral chromosomes. The evolution of chromosomes was characterized by two distinct episodes of whole-genome duplication (WGD). In accordance with this model, chromosomes Chr25 and Chr16 were fundamentally derived from a precursor chromosome owing to a recent WGD event, and they shared an older precursor chromosome with Chr5 and Chr6 prior to the ancient WGD.

The quality of the T2T genome was evaluated by comparison with the V3 assembly. In V4, the contig N50 showed marked improvement, rising from 465 kbp in V3 to 17.71 Mbp, a 38-fold increase. Furthermore, all 3,940 gaps that existed in V3 were eradicated in V4 (Supplementary Tables 5-6). The V4 genome is distinguished by its high continuity and coverage, as evidenced by the absence of any broken points or uncollapsed regions during the visualized assembly validation with ONT reads aligned onto chromosomes (Supplementary Fig. 4). The presence of high-copy rRNA and other regions with tandem repeats was confirmed by means of mapping depth. Upon analyzing the 4,146,102 bacterial artificial chromosome (BAC) sequences collected from the NCBI Nucleotide database, we mapped them onto the T2T genome. Our findings revealed that an astounding 95.62% of the sequences aligned concordantly with the genome, while the remaining sequences were predominantly highly repetitive, rendering them unmatched. Additionally, the mean RNA-seq mapping rates achieved were 96.74% and 96.52% for V4 and V3, respectively. Similarly, the DNA-seq mapping rates were 97.88% and 97.79% for V4 and V3, respectively. Notably, the LTR assembly index (LAI) values, which serve as a measure of assembly quality^25^, were 15.14 and 14.93 for V4 and V3, respectively (Supplementary Table 5 and Supplementary Fig. 1b). With a QV of 99.9986, the V4 consensus base quality demonstrates superior genome accuracy that exceeds Q40 and even achieves Q50. These assessments serve as evidence of the successful creation of a high-quality T2T genome.

### Synteny and Structural Variations in the V3 and V4 Genomes

The availability of the *P. patens* T2T genome presents an opportunity to conduct a more precise analysis of its structure. Our initial comparison using Mummer^26^ between the V4 and V3 versions revealed that most chromosomes maintained good collinearity (Fig. 1d). However, we observed large regions with disrupted synteny on Chr01, Chr16, Chr19, Chr22, and Chr23 in V3, with Chr16 exhibiting different orders and orientations at the chromosomal level (Supplementary Fig. 7a). This discrepancy may be attributed to the limited scaffolding ability of genetic linkage maps. To obtain a more realistic comparison between the two versions, we utilized V4 as a reference, fragmented V3 into contigs, reconstructed the V3 genome by RaGOO^27^, and identified structural differences between the two using SyRI^28^ (Supplementary Fig. 1d). A total of 253 contigs (totaling 227,515 bp in length) could not be integrated into the V4 version. These fragmented sequences comprise noncoding repetitive sequences. In comparison, the V4 genome shared 466 Mbp of syntenic regions with V3, which is 96.81% of the total genome size, leaving approximately 3.19% variant regions (Supplementary Fig. 1d). This indicates the existence of a discrepancy between the previous assembly of V3 and V4. As shown in Supplementary Table 7, 193 inversions (2.26 Mbp) and 322 translocations (3.67 Mbp) between V3 and V4 were identified, and 1.59 Mbp of V3 and 9.74 Mbp of V4 represented presence and absence variants (PAVs). However, the detected variations between the V3 and V4 genomes may have been overestimated, possibly due to differences in the assembly method and the comparisons caused by V3 breakpoints. The SyRI tool identified a total of 262,845 SNPs, while we utilized FreeBayes^29^ with V4 NovaSeq data for SNP calling in the V3 genome, which resulted in only 103,772 SNPs. The mean quantity of SNPs per 1 kbp is 0.22, indicating that there is no significant discrepancy between the genome sequencing materials obtained from the V3 and V4 versions. To accurately determine the pedigree of the *P. patens* employed in this research, we conducted an analysis using the protocol proposed by Haas et al^30^. By integrating Haas et al.’s SNP data with SNPs from the V4 genome, we constructed a neighbor-joining tree (Supplementary Fig. 2) for a total of 276 accessions. The results indicated that the *P. patens* presented in this study had the least SNP difference compared to Birmingham 2009 (1,238 SNPs, accounting for 0.84% of the total SNPs), which can be classified as the United Kingdom (UK) pedigree.

### Genomic Characterization and Annotation of V4 Genome

A total of 291 Mbp of repetitive sequences were detected in the V4 version, constituting 60.43% of the genome. Long terminal repeat retrotransposons (LTR-RTs) were the most prevalent repetitive sequences, accounting for 54.08% of the total. Among them, *Gypsy* elements were the most abundant, comprising 50.01% of the total, while *Copia* elements accounted for only 4.07% (Supplementary Table 8). Based on the estimation of insertion time using 11,968 noncentromeric LTR-RTs, a major burst occurred at 0.73 Mya during proliferation (Fig. 3e). Through the utilization of a significant RNA-seq dataset comprising 342 sample sets, we have effectively integrated two annotated versions, specifically V3 and NCBI Gnomon. After a manual correction process, we derived 29,836 protein-coding gene models that possess both start and stop codons (Supplementary Table 9). Among these models, 26,310 were supported by transcriptional evidence or had well-defined functional annotations, while 3,526 lacked such support. Notably, these unsupported genes were at least present in the annotated results from the V3 or NCBI version. Compared to the V3 version, which contained 32,458 protein-coding genes, the number of such genes was reduced by 2,622. This reduction was primarily attributed to the removal of TE-derived genes, the elimination of numerous duplicate genes (such as those with identical locations and duplicate names in the V3 version), and the exclusion of excessively short and incomplete gene sequences (Supplementary Table 10). We also identified 1,481 rRNAs, 641 tRNAs, and 524 noncoding RNAs (ncRNAs) (Supplementary Table 4). In contrast to the V3 version, which contained only 123 rRNA genes, the GBGF method successfully recovered the lost rRNA copies. Furthermore, utilizing Hi-C interactions that were sequenced by total DNA, we identified 1,212 topologically associated domains (TADs) that span across the chromosomes (Supplementary Table 11). These TADs serve as a means to visualize the chromosomal territories of all 26 chromosomes. As a result of its compact genome, the chromosomes of *P. patens* typically show a rosette configuration (Fig. 2b). In contrast to those of numerous flowering plants, the chromosomes of *P. patens* exhibit a more uniform distribution of both eu- and heterochromatin^14^. The confirmation of this perspective is evidenced by the distribution of A/B compartments in the V4 genome (Fig. 2a, Fig. 4a, Supplementary Figs. 12-37 and Supplementary Table 12).

**Fig. 3.**
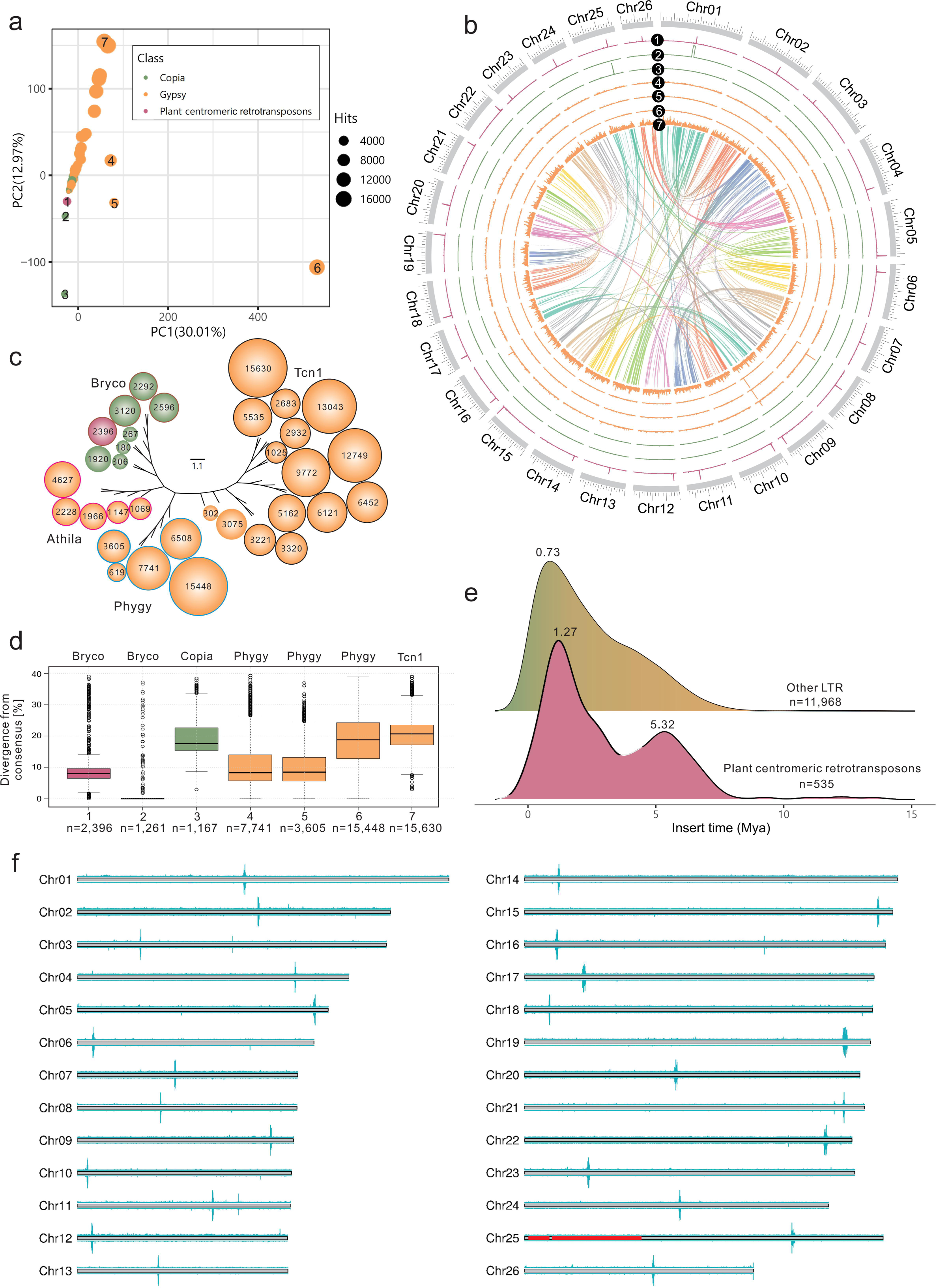
Characterization of centromeres. **a**, The distribution of LTR families in the genome as determined by the PCA method. The scatter plot displays each long terminal repeat (LTR) family as a distinct data point. The colors yellow, green, and purple represent Gypsy, Copia, and centromere-specific Copia, respectively. The size of each data point corresponds to the number of hits in the genome. The number of hits in the family is the primary determinant of the PC1 direction, while the PC2 direction is primarily influenced by the distribution characteristics. LTR families that exhibit a concentrated distribution in the region have a negative value in the PC2 direction. To provide a more detailed illustration of the distribution results, seven specific data points, labeled 1-7, were chosen. **b**, Circos plot demonstrating the distribution curves of the seven families along 26 chromosomes. **c**, A fruit tree exhibiting LTR family attributes, including evolutionary relationships and quantity. The nodes on the tree are distinguished by their assigned colors, which correspond to the specific LTR families to which they belong. The numerical values at the nodes indicate the number of families, while the colors at the edges of the nodes signify the subfamily classifications. **d**, Box plots for seven specific LTR families show the distribution of sequence divergence from their corresponding consensus sequences. Box edges represent the 0.25 and 0.75 quartiles, with the median values shown by bold lines. **e**, LTR insert time (Mya) for centromeric LTRs and other noncentromeric LTRs. The distribution curves have been marked with values indicating the peaks. **f**, ChIP-seq analysis from two independent lines. The top and bottom panels represent the results obtained using two independent CENH3-3×HA lines. In each chromosome, only one CENH3 peak was identified. The regions corresponding to the previously annotated Chr27 are shown in red.

**Fig. 4.**
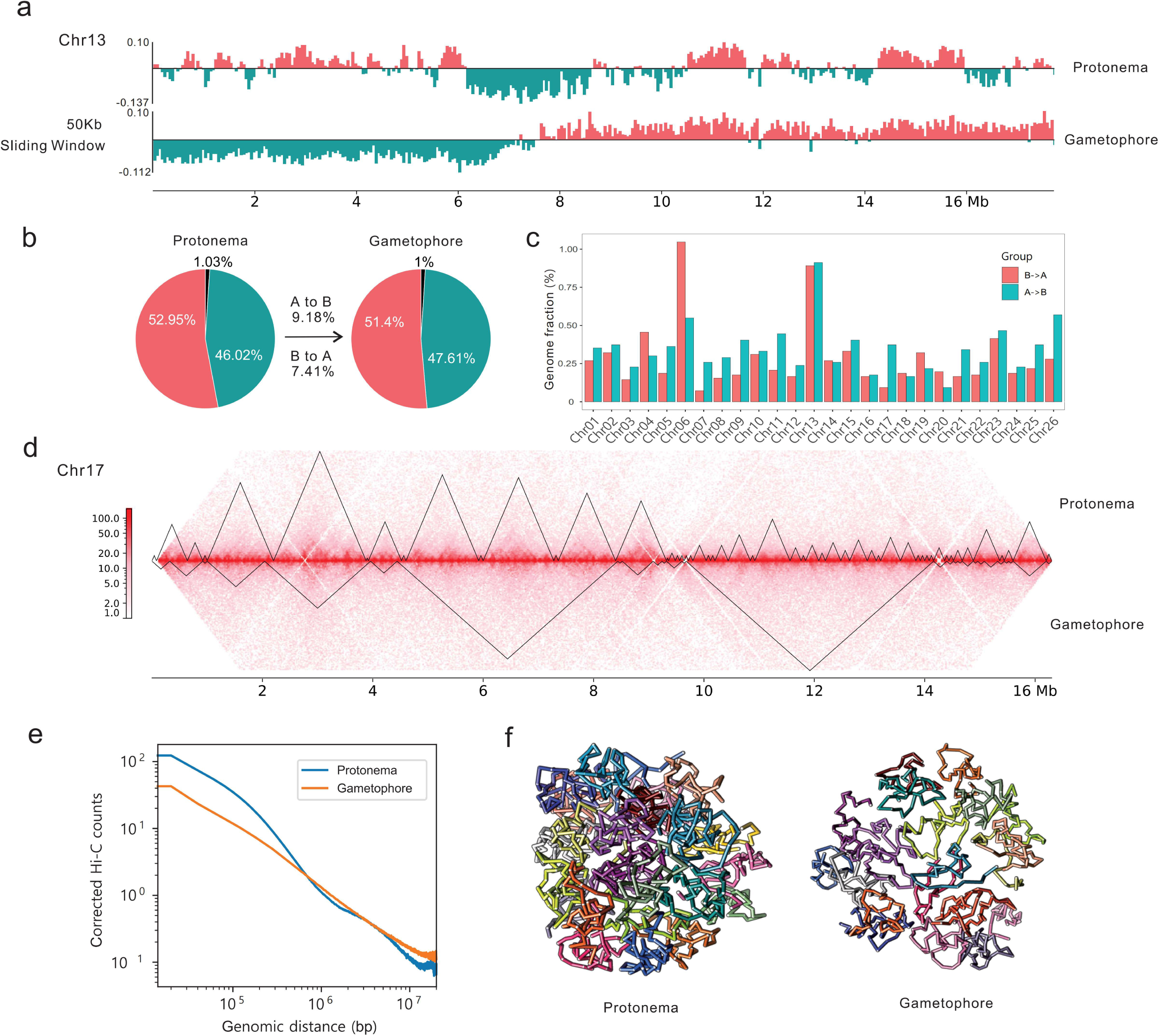
3D genome attributes of the protonema and gametophore. **a**, Alterations in A/B components of Chr13 under a 50 kbp sliding window. **b**, The pie charts show the proportion of the genome belonging to either the A (depicted in pink) or B (depicted in green) compartment in each of the two tissues that were analyzed. The regions with a principal component 1 (PC1) value of zero, which typically correspond to the highly repeated regions of the chromosomes, are represented in black. **c**, A bar chart depicting the proportion of genome differentiation in relation to A/B compartment switching across a total of 26 chromosomes. **d**, An example of the dynamic transformation of chromosome TAD boundaries is shown on Chr17. The Hi-C interaction heatmap shows the protonema and gametophore in the upper and lower portions, respectively. The triangular region represents the TAD interval that has been identified. **e**, Assessment of the attenuation in Hi-C interaction frequency as the interaction distance progressively expands. In comparison to the frequency of chromatin interaction observed in gametophores within a distance of 1 Mbp, the interaction intensity of the protonema exhibited a stronger trend as the distance decreased. This observation indicates that the protonema has a more condensed chromatin folding space. **f**, Whole-genome three-dimensional models for the protonema and gametophore. In contrast to the 3D arrangement of the protonema, the 3D organizational structure of the gametophore is less compact, featuring reduced interactions among intrachromosomal regions and a diminished number of TADs.

### Validation of 26 Chromosomes in V4 Genome

The structural differences between V4 and V3 are primarily characterized by two significant distinctions, one of which pertains to the variation in the number of chromosomes. In moss molecular genetics, as early as 1957, Bryan et al.^31^ suggested that the chromosome number was 27 based on microscopy observations, which was later supported by counting meiotic and mitotic chromosomes in 1993^32^. Since the inception of *P. patens* strain Gransden, it has been known that its chromosome number is 27. Subsequently, the draft genome and genetic linkage map of *P. patens* were successfully published in 2008^10^. In 2017, Lang et al.^14^ utilized BAC sequencing and a genetic linkage map to construct V3 chromosomes under the prior condition of n=27 (Supplementary Fig. 6). In contrast to the V3 version, the V4 version was obtained by employing a Hi-C structure-based approach to build chromosomes, which is a posterior conditional methodology that is not dependent on cytological observations. Additionally, early cytological observations were hindered by the condensed nature of the chromosomes, resulting in unclear images^31–33^. Thus, the key objective of this article is to settle the debate regarding the count of chromosomes. Chromosome-level synteny analysis showed that the region from the 5’ end to 5.4 Mbp of Chr25 in V4 was regarded as Chr27 of V3 (Supplementary Fig. 7a), resulting in the diversity of chromosome number between the two genomes. To elucidate this issue, we carried out conformational visualization by aligning our Hi-C data onto the three chromosomes. Our results, as presented in Supplementary Figure 7b, reveal that Chr25 and Chr27 in V3 show strong contact signals at each 5’ end. In contrast, Chr25 in V4 exhibits continuous interaction signals without any conformational errors. Simultaneously, we also authenticated the assembly of V4 Chr25, as depicted in Supplementary Figure 8. We effectively mapped reads exceeding 10 kbp to Chr25. ONT reads were consistently and uniformly aligned at the site where Chr25 and Chr27 were fused in V3, as well as its contiguous 5 kbp upstream and downstream regions, suggesting the absence of any assembly errors within this range. Moreover, in comparison to that in the V3 assembly, Chr25 in V4 has two telomeres located at both ends, unlike the solitary telomere present in both Chr25 and Chr27 in V3. These results suggest that Chr27 in V3 was incorrectly organized with a major inversion and was artificially separated as a distinct chromosome. This point is further substantiated by finding centromeres.

First, interchromosomal Hi-C interactions were examined and classified into two distinct categories based on their spatial orientation: telomeric and centromeric loci. Subsequently, a comprehensive Hi-C-based statistical analysis led to the identification of a total of 26 centromeric regions, each corresponding to one of the 26 chromosomes (Supplementary Fig. 9). Additionally, to elaborate on the features of these centromeres, we carried out a *de novo* exploration of these centromere regions. The centromeric and pericentromeric regions of chromosomes in higher plants are populated by *Gypsy* retrotransposons, giving rise to the chromovirus CRM clade^34^. Hence, through an examination of the distribution properties of these LTR elements, we can promptly identify the related LTRs and intervals that are specific to the centromere. With the help of TEsorter^35^, the 79 LTR families identified through RepeatModeler2^36^ were further categorized into subfamilies. The evolutionary relationships between these families were established by analyzing sequences with an integrase (INT) domain, with an integrity level above 75% (Fig. 3c). The distribution matrices for the aforementioned families were generated by employing a sliding window of 50 kbp across the genome. The principal component analysis (PCA) method was utilized to swiftly describe the distribution patterns of these matrices (Fig. 3a and Supplementary Table 13). The extent of dispersion manifested by the families in the PC1 and PC2 directions is influenced by the number of hits and the shape of the distribution, respectively. A distribution that is more concentrated is indicated by a lower PC2 value. As specifically stated, Fig. 3b presents the genome-wide distribution curves for the seven selected discrete points. Point 1 displays a distinct peak on each chromosome of V4, which corresponds to the plant centromeric retrotransposons (PCRs) of *P. patens*. Belonging to the Bryco clade within the *Copia* superfamily, this particular family is identified as the RLC5 element, as determined by Lang et al^14^. In comparison to the other five non-Bryco clade LTRs, the median of its internal divergence is at a lower level, indicating that the formation of the centromere in *P. patens* occurred during a relatively recent evolutionary period (Fig. 3d and Supplementary Table 14). The insertion time distribution of PCR exhibits two peaks, corresponding to 1.27 and 5.32 Mya (Fig. 3e), which aligns with the results of the similarity plot among 26 centromere regions (Supplementary Fig. 10). Finally, to validate the identification of the centromere regions in *P. patens*, we utilized the centromere-specific histone 3 variant (CENH3) and performed chromatin immunoprecipitation sequencing (ChIP-Seq). To do so, we replaced wild-type CENH3 with the fusion protein CENH3-3×HA through homologous recombination and obtained transgenic lines that were phenotypically similar to the wild type. We conducted ChIP using antibodies against HA in two independent lines and identified 26 peaks corresponding to the PCR-enriched regions of the 26 chromosomes in V4 by mapping reads associated with CENH3-3×HA obtained by ChIP (Fig. 3f). Both independent lines provided highly consistent results, as shown by the top and bottom peaks for each chromosome. We observed that regions corresponding to the previously annotated Chr27 lacked a CENH3 peak, providing additional evidence that this region is part of a chromosome. These centromere analyses provided strong support for the presence of only 26 chromosomes in *P. patens* and further suggested that the chromosomes can be classified into three types: metacentric, submetacentric, and subtelocentric chromosomes (Supplementary Table 15).

### Novel Telomere Structure and Updated Chromosomal Evolutionary Trajectory

Another significant structural deviation has been observed in the novel telomere. Specifically, out of the 52 telomeres present in the V4 assembly, 13 demonstrate a (GGCCCA)_n_ configuration (Fig. 2a and Supplementary Table 16). This novel configuration, which differs from the traditional (TTTAGGG)_n_ repeat, may be a secondary telomere structure resulting from the process of chromosome breakage and chromosome healing^37–39^ during evolution. The procedure aligns with the paleohistory of Chr12 and Chr26, which were believed to have emerged from the fragmentation of an ancient chromosome following the second whole-genome duplication (WGD) of *P. patens*^14^. Consequently, each of the two chromosomes is accompanied by a novel telomere situated at its terminus. Thus, the manifestation of the GGCCCA type potentially denotes chromosomal rupture, and this fracture signal cannot be inferred from the gene order collinearity of the extant genome. Drawing on this assumption, we have refined the paleohistory based on 26 modern chromosomes with such fracture signals. As illustrated in Fig. 2c, the gene collinearity of *P. patens* can be classified into seven clusters, each corresponding to one of the ancestral chromosomes. These chromosomes underwent two rounds of WGD during their evolutionary history. Our updated model, which deviates from Lang’s prior inference^14^, proposes that following WGD2, these chromosomes experienced 14 breaks, underwent 3 fusions, and lost 9 small fragments, ultimately giving rise to the current 26 chromosomes (Fig. 2d). In addition, it has been determined that the stability of centromere regions arose after the emergence of the 26 contemporary chromosomes. This is because chromosomes originating from a single disrupted ancestral intermediate possessed only one centromere. This outcome emphasizes the significant role played by LTRs in molding the structure of chromosomes throughout evolutionary processes. These findings pertaining to the telomere structure and the chromosomal evolutionary trajectory have shed light on the great efficiency of *P. patens* in rectifying double-strand breaks (DSBs) in DNA. When DNA DSB repair mechanisms fail to fix broken chromosomes, the process of chromosome healing is initiated, which culminates in the formation of new telomeres and the consequent emergence of new chromosomes^37–39^. This phenomenon can partially account for the diverse range of chromosome numbers observed in moss plants^31^.

### Dynamic Changes in 3D Genome Architecture between Protonema and Gametophore

By employing the T2T genome of *P. patens*, we can delve into the dynamic changes in 3D genome architecture during the developmental process or life cycle, thereby gaining a more profound understanding of the essential molecular mechanisms that govern the regulation of significant genes. In this study, we initiated a comparative study of the 3D genome structural differences between protonema and gametophore (Fig. 4). The transition from protonema to gametophore was a pivotal stage in the development of *P. patens*, facilitating its expansion from a two-dimensional to a three-dimensional plane. This phase is characterized by considerable spatial flexibility in the arrangement of the A/B compartments, with approximately 16.59% of the genome fraction changes occurring in these components (Fig. 4b, Supplementary Tables 17-18 and Supplementary Figs. 12-37). Notably, this alteration in A/B components is particularly marked in Chr6 and Chr13 (Fig. 4c). Previous studies have established that the genome can be categorized into two distinct regions, referred to as A and B, which contain relatively active and inactive regions, respectively. This A/B conversion may serve as the structural basis for expressing differences^40–41^. In comparison to the internal Hi-C interaction frequency of the protonema’s chromosome, the internal interaction frequency of the gametophore is lower, indicating a more relaxed chromatin structure. Additionally, the intensity of chromosomal interactions in the gametophore increased in comparison to that in the protonema (Supplementary Fig. 11). The findings of 3D structure fitting suggest that the gametophore genome has a looser structure, resulting in a reduction in the number of TAD domains from 1,318 to 998 (Fig. 4d, Supplementary Tables 19-20 and Supplementary Figs. 12-37). The findings indicate significant variations in the chromosome folding of protonema and gametophore, which can be linked to certain factors such as methylation levels and histone modifications (specifically H3K4me3, H3K27me3, and H3K9me2)^42–45^. Further exploration is essential to confirm and comprehend these factors as the principal determinants of genomic epigenetic traits.

## Conclusions

We have successfully constructed a T2T genome for *P. patens* ssp*. patens* (Gransden strain), which boasts exceptional contiguity and completeness. This achievement has rectified the previously held erroneous beliefs surrounding the chromosome count of *P. patens* and will serve as a valuable resource for in-depth genomic analysis and fundamental research on moss.

## Methods

### Sequencing material

*Physcomitrium patens* ssp*. patens* ecotype Gransden cultured at the Shenzhen Institutes of Advanced Technology, Chinese Academy of Sciences, was used as the sequencing material in the present study. *P. patens* was grown on BCDATG medium^46^ at 25LJ under 16 h light (30 μmol· m−2·s−1)/8 h dark conditions for 7 days. For DNA sequencing, 7-day-old protonemata were collected.

### DNA extraction and Illumina sequencing

The fresh materials were wrapped with filter paper to remove excess water and ground with a mortar and pestle in liquid nitrogen. Genomic DNA was prepared using the CTAB method. DNA purity was checked using a NanoPhotometer® spectrophotometer (IMPLEN, CA, USA), and DNA concentration was measured using a Qubit® DNA Assay Kit in a Qubit® 4.0 Fluorometer (Life Technologies, CA, USA). DNA integrity was checked by pulsed field gel electrophoresis (PFGE). A sequencing library was constructed with an insert length of 300 bp using the Illumina TruSeq^TM^ DNA Sample Preparation Kit (Illumina, San Diego, USA) following the manufacturer’s recommendations and sequenced on the MGISEQ-2000RS platform in a 150×150 bp paired-end run. In total, 51.14 Gbp of sequencing data were produced (Supplementary Table 1).

### Oxford Nanopore Technologies (ONT) library preparation and MinION sequencing

The purified DNA was used to prepare the long-read library (Oxford Nanopore MinION, UK) for third-generation genome sequencing. For MinION library construction, NEBNext Ultra II End-Repair/dAtailing Module (NEB_E7546) and NEBNext FFPE DNA Repair Mix (NEB M6630L) were used to perform DNA repair and end-prep. Adaptor ligation and clean-up of the large DNA fragments were performed by a Ligation Sequencing Kit (SQK-LSK109). A Flow Cell Priming Kit (EXP-FLP001) was used to prepare the priming buffer to prime the MinION flow cells (FLOMIN106D R9.4.1 version). Sequencing began immediately in the MinION^TM^ portable single-molecule nanopore sequencer (Oxford Nanopore Technologies Limited, UK) using MinKNOW 3.1.8 software (Oxford Nanopore Technologies Limited, UK). The read event data were base-called by Oxford Nanopore base caller Guppy version 2.1.3 with fast-base calling settings. A total of 42.98 Gbp of reads were obtained (Supplementary Table 1). The mean and N50 length of all reads reached 8.21 and 31.63 kbp, respectively.

### Hi-C sequencing

To construct the Hi-C library, frozen *P. patens* materials were thawed slowly on ice and suspended in 45 ml of 37% formaldehyde in serum-free Dulbecco’s modified Eagle’s medium for chromatin cross-linking. After incubation at room temperature for 5 min, glycine was added to quench the formaldehyde, followed by incubation at room temperature for another 5 min and then on ice for over 15 min. The cells were further lysed in prechilled lysis buffer (10 mM NaCl, 0.2% IGEPAL CA-630, 10 mM Tris-HCl, and 1× protease inhibitor solution) using a Dounce homogenizer. The chromatin was digested by MboI restriction enzyme and labeled with biotin, followed by ligation. The Hi-C library was sequenced on a NovaSeq 6000 platform (Illumina, USA) and generated reads with a length of 150 bp producing 32.08 Gbp of data in total (Supplementary Table 1). To contrast the 3D genome structure variances between protonema and gametophore, we carried out Hi-C library construction and sequencing on both tissues separately, resulting in the generation of 36 Gbp and 43 Gbp of data for protonema and gametophore, respectively (Supplementary Table 1).

### K-mer analysis and genome size estimation

A k-mer approach was utilized to determine the size of the haploid genome, potential heterozygosity, and repeat length of the genome via analysis of the DNA sequencing library. The raw data were processed through fastp v0.20.1^47^ to eliminate any adaptors, low-quality bases, and reads that contained more than 10% unknown bases (“N”). GenomeScope v1.0^48^ was then employed to estimate the assembly based on histograms created from the resulting 17-, 19-, 21-, and 23-mers using Jellyfish v2.3.0^49^. The estimated genome depth for 21-mers was utilized to determine the seed cutoff needed for NextDenovo v2.3.0^22^ (https://github.com/Nextomics/NextDenovo). The genome size of *P. patens* was estimated to be between 460.24 and 480.15 Mbp, and the heterozygosity was determined to be between 0.145 and 0.166%, indicating that a haploid genome was present (Supplementary Fig. 1a).

### Identification of the pedigree of the *P. patens* accession sequenced in this study

To precisely ascertain the pedigree of the genome sequencing material in this study, it was essential to follow the method delineated by Haas et al^30^. The SNP data obtained from 275 accessions by Haas et al, in conjunction with the SNPs between V3 and V4, were included in this process. Specifically, we employed the V3 genome as the reference and scrutinized the Illumina reads derived from the genomic DNA of V4. We performed SNP calling using Freebayes^29^ and subsequently constructed the consensus sequence utilizing BCFtools^50^. From the results of Haas et al., we extracted the SNP sites and created an SNP fasta file for V4. By utilizing the resultant 276 copies of the 146,816 SNP matrix, a neighbor-joining tree (with 100 replicates) was generated using TreeBeST^51^ to deduce the lineage of V4 (Supplementary Fig. 2).

### Preliminary assembly, chromosome construction and gap filling with a graph-based gap filling (GBGF) approach

Prior to the formal assembly, we utilized NextDenovo^22^, Flye^23^, and NECAT^52^ as assemblers or read calibration tools. To evaluate discrepancies in the results among distinct datasets and assemblers to select the best genome backbone for gap filling, we tested six different combinations, incorporating all ONT reads or filtering out the organelle read dataset (Supplementary Table 2). ONT reads derived from organelle genomes were removed by the aligner Minimap2 v2.17-r941^53^. Before alignment, considering a circular topology, the heads and tails of the chloroplast (accession No. NC_005087) and mitochondrial genomes (accession No. KY126309) were extended by the corresponding 50 kbp. Following an exhaustive evaluation procedure, it was ascertained that assembled contigs from NextDenovo, which made use of all obtainable data, were the best. This preliminary assembly had a total length of 473.26 Mbp (contig N50: 16.62 Mbp) spread across 45 contigs. This contig assembly was named s1. Raw Hi-C sequencing reads were first trimmed by fastp v0.20.1^47^. To remove the invalid pairs of Hi-C reads without effective ligation, the reads were mapped to the s1 assembly using Bowtie2 version 2.3.5.1^54^, and effective Hi-C contact reads were retrieved by bamTofastq (https://github.com/10XGenomics/bamtofastq) based on the final BAM file generated by HiC-Pro version 2.11.1^55^. Juicer version 1.5^56^ was used to align filtered Hi-C reads to the s1 assembly. The resultant binary contact matrix was used for manual contig scaffolding using the 3D *de novo* assembly (3D-DNA) pipeline version 180114^57^. Misassemblies of contigs were manually corrected based on the Hi-C contact maps. Only 26 chromosome frames were acquired. This scaffolding assembly was named s2. Out of the 26 chromosomes, 10 are devoid of gaps, while the remaining chromosomes have a cumulative total of 23 gaps. Furthermore, a total of 47 telomere structures were detected at both ends of these chromosomes, encompassing not only the conventional (TTTAGGG)_n_ configuration but also the (GGCCCA)_n_ configuration. Next, the graph-based gap filling (GBGF) approach, a method devised by our team, was utilized to perform gap filling on the s2 assembly.

GBGF is a partially manual method that seeks to investigate the factors that contribute to the occurrence of genomic breakpoints. Subsequently, it reconstructs the path of breakpoint positions on the assembly graph to ensure gap filling. To summarize, the formation of breakpoints can be categorized into simple breaks that lack read coverage, breaks stemming from heterozygous intervals in diploids, local assembly errors in sequences, and certain unique structures such as highly tandem repeats, reverse repeats, and complex regions that involve the cross-linking of these repeating fragments (Fig. 1b, Supplementary Fig. 3). The GFA (Graph Fragment Assembly) of the unitig graph represents the smallest unit that forms a contig and serves as the prefile for the GBGF method. The utilization of Flye^23^ facilitates the creation of GFA files. The utilization of distinct read correction methods and datasets can influence the construction outcomes of GFA. Therefore, we developed five GFA graphs utilizing various read correction data obtained from the initial genome assembly (Supplementary Table 3).

Afterward, it is necessary to anchor these gaps onto the graphs. Specifically, 50-200 kbp upstream and downstream sequences around a gap or a region were extracted and anchored to corresponding edges through BLASTN v2.2.31^58^ (Evalue: 1e-100). Then, Bandage v0.7.1^59^ was employed to visualize and explore the reasonable path between anchors. Concurrently, it is essential to factor in the sequencing depth particulars of these edges to ascertain the exact number of copies accurately. In instances where interval paths are complex, we employed minigraph^60^ to map ONT reads on these graphs. This enables us to identify reads that can span these intervals and aid in the construction of the appropriate path. The resultant sequence derived from this path can then be utilized to replace the gap interval and effectively complete the filling process. Genome assembly may face several challenges leading to breakpoint formation, including structural conflicts arising from terminal redundancy and complex structures such as reverse duplications. While overlap-based gap-filling software may prove useful, it is not always efficient in completing the process. GBGF may not be the most efficient method, but it certainly proves effective in such cases. Additionally, the overlap-based approach may overlook the existence of concatenated duplicates because it fails to consider the corresponding number of copies within the interval. By employing GBGF, we filled the 23 gaps present in the s2 genome, yielding a raw genome without gaps.

When confronted with chromosomes that lack telomere sequences, our primary strategy involves utilizing the GBGF method to identify and introduce these sequences. If this approach fails, we will then resort to searching for and integrating corrected long reads to compensate for the absence of telomere sequences. After polishing by corrected ONT reads, another three rounds of polishing were performed using Hapo-G^61^ with Illumina short reads to improve the accuracy of the T2T genome.

The final T2T genome was named version V4. It contains a total length of 481.75 Mbp with the same contig N50 and scaffold N50 (17.71 Mb) spread across 26 chromosomes and 52 observed telomeres. An overview of the whole genome assembly pipeline is shown in Fig. 1. Heatmap visualization of genome-wide Hi-C interactions was generated by Juicerbox v1.11.8 (https://github.com/aidenlab/Juicebox) to evaluate contact maps of the V4 assembly (Supplementary Fig. 5).

### Evaluation of genome assembly quality

#### 1) Mapping of RNA reads against the assembly

The completeness of coding potential in two *P. patens* assemblies, namely, V4 (this study) and V3, was evaluated using RNA-seq reads (342 samples) obtained from the NCBI SRA database (Supplementary Table 21). Prior to using HiSAT2 version 2.1.0^62^, all cDNA libraries were combined. The overall mapping rates for V4 and V3 were 96.60% and 96.52%, respectively. As illustrated above, the mapping rate to V4 was higher, although no significant variation was noted.

#### 2) Mapping of DNA reads against the assembly

To assess the quality of the genome assembly, we utilized the paired-end (PE) library with an insert size of 300 bp from the present study to examine read mapping rates. Using BWA version 0.7.12-r1039^63^ with default parameters, we mapped all reads back to the assembled genome and subsequently evaluated the overall mapping rate using SAMtools version 1.9^64^. The mapping rates for V4 and V3 were 97.88% and 97.79%, respectively. The mapping rate for V4 was found to be higher, particularly in terms of properly mapped read pairs, which reached 97.88% (mapped) and 97.11% (properly paired), as anticipated.

#### 3) Consensus base quality, LTR assembly index and assembly integrity

The consensus base quality of the V4 assembly was determined using the POLCA pipeline^65^, which is available in MaSuRCA version 3.4.1^66^. The resulting quality value (QV) was 99.9989, indicating high genome accuracy above Q40. To evaluate genome contiguity based on predicted intact LTRs, the LTR assembly index (LAI) was used as a novel indicator provided by LTR_retriever^25^. The LAI values for V4 and V3 were 15.14 and 14.93, respectively, and were positively correlated with the quality of the assembly. To validate and visualize the V4 assembly, Tapestry^67^ was employed to align nanopore long reads with lengths above 10 Kbp.

#### 4) BUSCO evaluation

We performed Benchmarking Universal Single-Copy Orthologs (BUSCO)^68^ analysis using the embryophyta_odb10 dataset on two existing *P. patens* assemblies (V4 and V3) (Supplementary Table 5). Our findings revealed that both assemblies captured the same percentage (88.1%) of complete BUSCOs, indicating comparable completeness levels.

### Transposable element annotation

For homology-based prediction, we used RepeatMasker v4.0.7^69^ to search against Repbase^70^. For *ab initio* prediction, we used Tandem Repeats Finder v4.07b^71^, RepeatModeler v1.0.8 (http://www.repeatmasker.org/RepeatModeler/), Piler v1.0^72^ and RepeatScout v1.0.5^73^ with default parameters. Full-length long terminal repeat retrotransposons (LTR-RTs) were identified using LTR_finder v1.07^74^ and LTRharvest^75^. Redundancies in the full-length LTR-RTs were then removed using LTR_retriver^25^. The substitution rate between the two end sequences of each LTR-RT was calculated using PAML^76^. LTR insertion time was estimated according to the formula T=S/2μ, where S is the substitution rate and μ is the mutation rate (9 E−9 per site per year)^77^.

Overall, the identified repeat sequences in the V4 genome accounted for 60.10% of the total genome length, which was 286.55 Mbp (Supplementary Table 8). Long terminal repeats (LTRs) were the most abundant, comprising 53.75% of the genome. We identified 3,309 intact LTR-RTs in the V4 genome. The estimation of insertion time for 11,968 LTR elements revealed that one burst took place 0.73 million years ago (Mya) (Fig. 3e). The same approach was employed to examine the insertion time of 535 LTRs that are associated with centromeres (Fig. 3e).

### Whole-genome duplication and updated karyotype evolutionary scenario of *P. patens*

Whole-genome duplication (WGD) analysis was performed by searching for collinearity within the V4 genome using wgdi^78^. Duplicated gene pairs in collinear blocks were extracted for sequence alignments. For each pair, the nonsynonymous (*Ka*) and synonymous substitution (*Ks*) rates were calculated using the yn00 method implemented in PAML^76^. We obtained anchor genes from the intragenomic synteny blocks present in a ratio of 1:3. An obviously discrepant *Ks* distribution according to the color scale (Fig. 2c) revealed two rounds of WGD events. The karyotype evolutionary model of chromosomes has been augmented with a novel break signal (GGCCCA telomere) in the Lang inference result. Additionally, a new fusion event was identified through collinearity analysis, which is represented as the h+l event in Figure 2d.

### A/B compartments, topologically associating domains (TADs) and 3D genome model

TADs were called using Armatus v.2.3^79^, setting the gamma-max parameter to 0.6 at a 40 kbp resolution. For A/B compartment profiles, we used the Juicer Tool from the Juicer package to compute the first eigenvector of each chromosome from global contact matrices. The correlation coefficient between gene density and the eigenvector was computed, and B compartments were assigned negative values of the eigenvector that corresponded to regions with lower gene density. PyGenomeTracks^80^ was employed to visualize A/B department profiles and TAD boundaries for each chromosome. The hicPlotDistVsCounts function within HiCExplorer^81^ was used to generate the plot that displays the relationship between distance and Hi-C counts. The Hi-C chromatin interaction z score analysis (protonema/gametophore and gametophore/protonema) at a 500 kbp resolution was performed by Juicerbox v1.11.8.

Chrom3D v1.0.2^82^ was used to generate the three-dimensional genome model. Using predetermined TAD regions, Chrom3D constructed a GTrack file that recorded the relative interaction strength among TADs. In the final results, each TAD is represented by a bead, the size of which is proportional to the span size. The final model was generated using the following settings: Chrom3D -l 10000 -y 0.15 -r 5.0 -n 3000000. Three-dimensional models were visualized by Chimera v1.16^83^.

### Genome assembly comparison

To assess the quality of our assembled genome, we first compared genome-wide alignments of V4 and V3 performed by Mummer v3.23^26^ with the command “nucmer --maxmatch -c 500 -b 500 -l 1000” and the parameter “delta-filter -m -i 90 -l 100”, which resulted in the best alignments with at least 100 bp matches and at least 90% identity between the two assembled genomes. A synteny dotplot was drawn using an online website (https://dot.sandbox.bio/) (Supplementary Fig. 7a). Due to the limitations of the scaffolding ability employed by the V3 genome, numerous structural variations might have been inaccurately identified as false positives. To ensure precise classification and statistical analysis of such variations, we used V4 as a reference and fragmented V3 into contigs. These contigs were then reconstructed into the V3 genome using RaGOO to ensure accuracy. Mummer alignments were performed by comparison between V4 and this new V3. The alignment results were then used as input in SyRI v1.2^28^ to discover genome duplications, translocations, inversions and syntenic, unaligned, and divergent sequences (Supplementary Table 7). Considering that both the V4 and V3 assemblies were sequenced from the same *P. patens* strain, large structural variations, mainly including inversions and translocations, were unexpected. Thus, to validate the structural differences between the V4 and V3 genomes, Hi-C reads sequenced in this study were also aligned to the original V3 genome through the Juicer pipeline. A heatmap displaying the structural conformation of Chr25 and Chr27 in V3 was visualized by Juicerbox v1.11.8 (Supplementary Fig. 5). Notably, two chromosomes (Chr25 and Chr27) in the V3 genome are composed of one chromosome corresponding to Chr25 in V4.

### Gene prediction and annotation

Protein-coding genes were retrieved using EVidenceModeler (EVM)^84^ by integrating transcript evidence, *ab initio* prediction and protein homology searching. For transcript evidence, RNA-seq reads from large datasets (342 samples) (Supplementary Table 21) were aligned against the V4 genome and assembled into transcripts using StringTie v2.1.5^85^. *De novo* transcriptome assembly was performed by SPAdes v3.14.0^86^ with the parameter --rna. Combined transcripts were used to assist genome annotation by aligning them to the genome using PASA^84^. The resultant high-confidence gene model sets that shared a length above 900 bp as well as start and stop codons were used as training datasets in AUGUSTUS v3.4.0^87^ and SNAP v2006-07-28^88^. For *ab initio* prediction, AUGUSTUS, SNAP, and Genemark-ES v4.48_3.60_lic^89^ were employed. For protein homology searching, protein sequences from two open-access *P. patens* annotations, including the NCBI assembly database and PETAmoss database, were collected and aligned to the V4 genome using GeMoMa^90^. Consensus gene models generated from EVM were further updated by PASA in two rounds. Following the filtration of gene models derived from transposable element coding domains (e.g., PF03732: Retrotrans gag) and the subsequent manual correction of each individual model, a total of 29,836 gene models were ultimately obtained (Supplementary Table 4). Genes that possess well-defined functional annotations and are substantiated by transcriptome evidence are categorized as high-quality (HQ) genes. Conversely, genes that do not exhibit such evidence must exhibit a minimum of two overlaps in three annotations (NCBI version, V3 version, and *ab initio* prediction) to be preserved. These genes are classified as low-quality (LQ) genes. Compared to the previous version (V3), which had a total of 32,458 genes that code for proteins, the subsequent iteration exhibited a decrease of 2,622 such genes. This reduction was primarily attributed to the elimination of genes that originated from transposable elements, the removal of numerous duplicate genes (including those with identical gene locations and duplicate gene names in the V3 version), and the exclusion of gene sequences that were excessively short or incomplete. Supplementary Table 10 presents the association between the V4 genes and V3 genes. Additionally, 423 novel gene loci were integrated.

For annotation, predicted protein-coding genes were searched against the nr^91^, Swiss-Prot^92^, KEGG^93^, KoFamKoala^94^, eggNOG^95^, InterPro^96^, Pfam^97^, CDD^98^ and PANNZER2^99^ databases (Supplementary Table 9). GO term annotation was carried out by integration of GO hits from PANNZER2 and Blast2GO^100^. rRNA was identified by barrnap v0.9 (https://github.com/HRGV/barrnap). tRNAScan-SE v2.0.7^101^ was used with default parameters to identify tRNA genes. In addition, miRNAs and snRNAs were identified by mapping the genome sequences to the Rfam database (http://rfam.xfam.org/) using INFERNAL^102^.

### Identification of centromere locations

In addition to the true spatial proximity among loci resulting in long-range interactive hotspots in the Hi-C heatmap, Hi-C contacts among regions enriched in similar elements will have more signals than those of unique sequences. Thus, hotspots of interchromosomal Hi-C interactions could favor the identification of constitutive elements such as telomeres and centromeres. To elucidate this phenomenon, a heatmap of interchromosomal Hi-C interactions was created (Supplementary Fig. 9). Two types of obvious hotspots could be observed visually. Hotspots located at the borders of chromosomes were telomere interactive loci, while the remaining hotspots highlighted inside chromosomes were centromere interactive loci.

To capture centromere-like regions, we first created 50 kbp nonoverlapping windows across chromosomes using BEDTools v2.30.0^103^. Then, we filtered significant window-window interactions using the FDR (cut off = 0.01) calculated by the noncentral hypergeometric (NCHG) test^104^. We further removed filtered windows that existed at both ends of the chromosome. As these filtered regions were noncoding regions with similar element components, we also ran a principal component analysis (PCA) through a matrix built by calculating the numbers of each predefined LTR family in every preassigned window to identify the LTR families underlying the composition of centromeres (Fig. 3a). The evolutionary relationships between these families were determined utilizing the neighbor-joining tree in MEGA5^105^. This involved scrutinizing sequences that possessed an integrase (INT) domain and had an integrity rating surpassing 75% (Fig. 3c). We discovered a *Copia* family related to the centromere belonging to the Bryco clade. This particular family is characterized by a centralized distribution pattern that is enriched in windows overlaid with NCHG results. Furthermore, each of these windows exhibits one peak on each chromosome. Similarity plots among 26 centromere regions were made by StainedGlass^106^.

### Plasmid construction and *Physcomitrella patens* transformation

To obtain CENH3-3 × HA transgenic lines, two 1 kbp regions flanking the CENH3 (Pp01Ghq00749) stop codon were cloned into BJ36 as homologous arms. Then, 3×HA and G418 resistance genes were added between the two homologous arms. The plasmid was linearized by NotI and transformed into *Physcomitrium patens* by PEG^107^. Several transgenic lines were obtained after G418 resistance screening, and two of them were used in the following experiments.

### Chromatin immunoprecipitation

Chromatin immunoprecipitation (ChIP) was performed as previously described with minor modifications^108^. Seven-day-old protonema of two CENH3-3×HA lines were collected and fixed. After chromatin extraction and sonication, chromatin was bound to the anti-HA antibody (Roche, Catalog Number: 11583816001) with Dynabeads Protein A (Invitrogen, Catalog Number: 10002D). The purified DNA was amplified for high-throughput sequencing by an Ultra II DNA Library Prep Kit (New England Biolabs, Catalog Number: E7645S). Then, the libraries were sequenced, and the 150-bp reads were analyzed by Trimmomatic^109^ and BWA^63^. The analysis results were obtained based on the V4 genome by MACS2^110^ and drawn by karyoploteR^111^. We estimated the precise locations of centromeres based on these peak regions (Supplementary Table 15).

## Acknowledgments

We thank Dr. Chunli Chen from Huazhong Agricultural University and Ms. Changxiu Yu from the Agricultural Genomics Institute at Shenzhen for their advice on chromosome analysis. We also thank Prof. Haodong Chen at Tsinghua University for providing assistance with *P. patens* genetics.

## Authors’ contributions

J.Y.,Y.M., and J.D. conceived the study. J.Y. and J.D. managed the major scientific objectives. J.Y. and G.B. designed the T2T genome assembly, evaluation, and data analysis. J.W.Y. managed the plant materials. H.W. helped with the assembly and annotation. S.Z., M.Z., Y.S., and X.H. collected the sequenced samples. J.W.Y., Y.J. Y.M., and J.D. designed and performed chromatin immunoprecipitation. J.Y. and G.B. led the article preparation, together with J.W.Y., H.W. and J.D. All authors read and approved the final article.

## Availability of data and materials

The genome assembly and annotations have been deposited in Figshare with DOI: doi.org/10.6084/m9.figshare.22975925.v1. The Illumina reads (genomic sequencing and Hi-C) and Nanopore reads generated in this study were deposited into the NCBI SRA with the BioProject ID PRJNA742485.

