## Supplementary Information for "Telomere-to-telomere genome of the model plant *Physcomitrium patens*"

***This file includes:***

**Supplementary Figures**

**Supplementary Figure 1.** A fundamental overview of the main content presented in this work.

**Supplementary Figure 2.** The neighbor-joining cladogram tree of five *P. patens* accessions built by SNPs derived from Haas et al. (2020).

**Supplementary Figure 3.** The process of graph-based gap-filling for the remaining 16 gaps.

**Supplementary Figure 4.** Genome assembly validation achieved by analyzing sequencing coverage and depth in relation to the 26 chromosomes in *P. patens*.

**Supplementary Figure 5.** Hi-C interactions among 26 chromosomes at a 500 kbp resolution.

**Supplementary Figure 6.** A compilation of pertinent time nodes in the study of chromosome numbers as well as notable research papers that explore the advancements in the field of *P. patens* genome research.

**Supplementary Figure 7**. Structural conflicts between two distinct versions of particular chromosomes.

**Supplementary Figure 8.** Assembly accuracy validation for Chr25 in V4 by read mapping.

**Supplementary Figure 9.** Interchromosomal Hi-C interactions exhibit two types of hotspots according to the positions on each chromosome.

**Supplementary Figure 10.** Dot plot for centromere intervals of 26 chromosomes along with their respective upstream and downstream 20 kbp intervals.

**Supplementary Figure 11.** The Hi-C chromatin interaction z score (protonema/gametophore and gametophore/protonema) at a 500 kbp resolution.

**Supplementary Figure 12.** A/B compartment switching and the alteration of topologically associating domain (TAD) boundaries between protonema and gametophore on Chr01.

**Supplementary Figure 13.** A/B compartment switching and the alteration of topologically associating domain (TAD) boundaries between protonema and gametophore on Chr02.

**Supplementary Figure 14.** A/B compartment switching and the alteration of topologically associating domain (TAD) boundaries between protonema and gametophore on Chr03.

**Supplementary Figure 15.** A/B compartment switching and the alteration of topologically associating domain (TAD) boundaries between protonema and gametophore on Chr04.

**Supplementary Figure 16.** A/B compartment switching and the alteration of topologically associating domain (TAD) boundaries between protonema and gametophore on Chr05.

**Supplementary Figure 17.** A/B compartment switching and the alteration of topologically associating domain (TAD) boundaries between protonema and gametophore on Chr06.

**Supplementary Figure 18.** A/B compartment switching and the alteration of topologically associating domain (TAD) boundaries between protonema and gametophore on Chr07.

**Supplementary Figure 19.** A/B compartment switching and the alteration of topologically associating domain (TAD) boundaries between protonema and gametophore on Chr08.

**Supplementary Figure 20.** A/B compartment switching and the alteration of topologically associating domain (TAD) boundaries between protonema and gametophore on Chr09.

**Supplementary Figure 21.** A/B compartment switching and the alteration of topologically associating domain (TAD) boundaries between protonema and gametophore on Chr10.

**Supplementary Figure 22.** A/B compartment switching and the alteration of topologically associating domain (TAD) boundaries between protonema and gametophore on Chr11.

**Supplementary Figure 23.** A/B compartment switching and the alteration of topologically associating domain (TAD) boundaries between protonema and gametophore on Chr12.

**Supplementary Figure 24.** A/B compartment switching and the alteration of topologically associating domain (TAD) boundaries between protonema and gametophore on Chr13.

**Supplementary Figure 25.** A/B compartment switching and the alteration of topologically associating domain (TAD) boundaries between protonema and gametophore on Chr14.

**Supplementary Figure 26.** A/B compartment switching and the alteration of topologically associating domain (TAD) boundaries between protonema and gametophore on Chr15.

**Supplementary Figure 27.** A/B compartment switching and the alteration of topologically associating domain (TAD) boundaries between protonema and gametophore on Chr16.

**Supplementary Figure 28.** A/B compartment switching and the alteration of topologically associating domain (TAD) boundaries between protonema and gametophore on Chr17.

**Supplementary Figure 29.** A/B compartment switching and the alteration of topologically associating domain (TAD) boundaries between protonema and gametophore on Chr18.

**Supplementary Figure 30.** A/B compartment switching and the alteration of topologically associating domain (TAD) boundaries between protonema and gametophore on Chr19.

**Supplementary Figure 31.** A/B compartment switching and the alteration of topologically associating domain (TAD) boundaries between protonema and gametophore on Chr20.

**Supplementary Figure 32.** A/B compartment switching and the alteration of topologically associating domain (TAD) boundaries between protonema and gametophore on Chr21.

**Supplementary Figure 33.** A/B compartment switching and the alteration of topologically associating domain (TAD) boundaries between protonema and gametophore on Chr22.

**Supplementary Figure 34.** A/B compartment switching and the alteration of topologically associating domain (TAD) boundaries between protonema and gametophore on Chr23.

**Supplementary Figure 35.** A/B compartment switching and the alteration of topologically associating domain (TAD) boundaries between protonema and gametophore on Chr24.

**Supplementary Figure 36.** A/B compartment switching and the alteration of topologically associating domain (TAD) boundaries between protonema and gametophore on Chr25.

**Supplementary Figure 37.** A/B compartment switching and the alteration of topologically associating domain (TAD) boundaries between protonema and gametophore on Chr26.

**Supplementary Tables in a separate Excel file**

**Supplementary Table 1.** Statistics of whole-genome sequencing data.

**Supplementary Table 2.** Statistics of six genome assemblies with different dataset combinations and software.

**Supplementary Table 3**. The data sources for GFA (graph fragment assembly) files and the acquisition of graph statistics.

**Supplementary Table 4**. Summary statistics of the V4 genome assembly.

**Supplementary Table 5**. Completeness comparison between V4 and V3.

**Supplementary Table 6**. Length and gap number comparison between V4 and V3.

**Supplementary Table 7**. Summary statistics of SyRI analysis.

**Supplementary Table 8**. Transposable elements predicted from the V4 genome.

**Supplementary Table 9**. Summary statistics of gene annotations.

**Supplementary Table 10**. Corresponding relationship between the V4 and V3 versions of the gene.

**Supplementary Table 11**. A total of 1,212 TAD intervals identified through Hi-C sequencing of total DNA.

**Supplementary Table 12**. Distribution of A/B compartment intervals across 26 chromosomes.

**Supplementary Table 13**. Information on the 79 families that were involved in the process of PCA.

**Supplementary Table 14**. Box plot statistics of seven selected data points regarding divergence.

**Supplementary Table 15**. Centromere regions in the V4 genome.

**Supplementary Table 16**. Telomere sequences in the V4 genome.

**Supplementary Table 17**. Distribution of A/B compartment intervals across 26 chromosomes in the protonema.

**Supplementary Table 18**. Distribution of A/B compartment intervals across 26 chromosomes in the gametophore.

**Supplementary Table 19**. Identified 1,318 TAD intervals in the protonema.

**Supplementary Table 20**. Identified 996 TAD intervals in the gametophore.

**Supplementary Table 21**. RNA-seq datasets used in gene prediction.

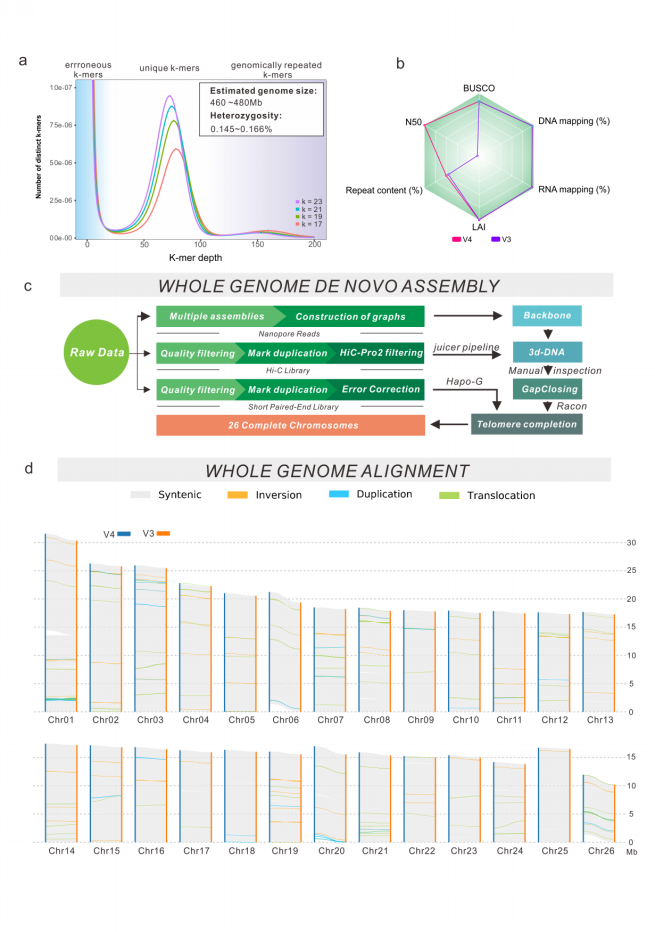
**Supplementary Figures**

**Supplementary Figure 1. A fundamental overview of the main content presented in this work**. **a**, Distribution of 17-23 K-mer frequencies in the *P. patens* genome. **b**, A radar chart was utilized to show the quality disparity between the V4 genome and its antecedent, V3. The evaluation was based on six distinct indicators, and the findings were scrutinized to identify any discrepancies in quality between the two versions. **c**, A concise diagram illustrating the process of V4 assembly. d, Results of SyRI analysis showing genome sequence collinearity and structural variants. To ensure the utmost precision in capturing the genuine discrepancies between the two genome versions, the V3 sequence was fragmented into contigs (where N bases were interrupted). Then, using RaGOO software, 26 pseudochromosomes were created to align with V4.

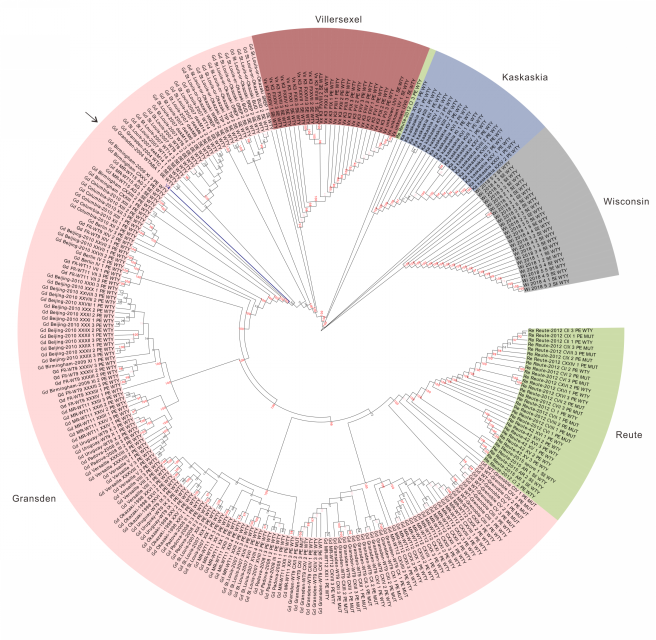

**Supplementary Figure 2. The neighbor-joining cladogram tree of five *P. patens* accessions built by SNPs derived from Haas et al. (2020).** The genome sequencing material used in this study is denoted on the tree by an arrow. Bootstrap values under 100 replicates are shown on nodes.

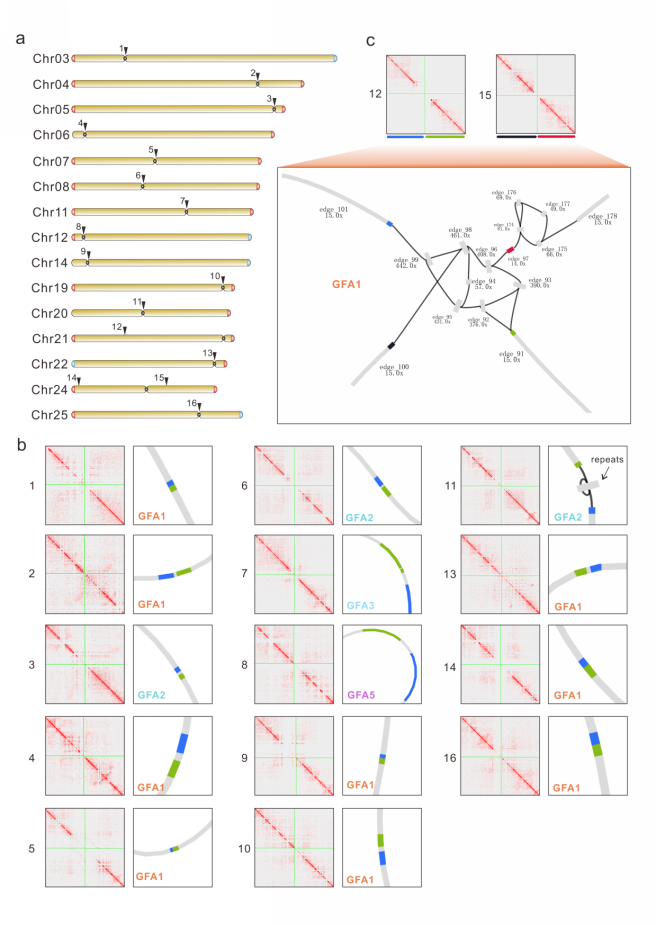

**Supplementary Figure 3. The process of graph-based gap-filling for the remaining 16 gaps**. **a**, Fifteen chromosome karyotype maps exhibit the remaining 16 gaps, which are marked with numerical values. The position of the centromere is the region where the chromosome constriction is situated. **b**, Locations of breakpoints (or gaps) that are anchored to the corresponding graphs. The Hi-C heatmaps accurately depict gap intervals with a resolution of 1 kbp, along with the identification of the breakpoints in the corresponding graph. On a corresponding edge, the precise locations of the 14 intervals where gaps occur on the graph are marked (blue and green bands), thereby enabling effortless gap-filling. **c**, Two gaps occur in the complex region. The upstream and downstream intervals of two gaps are labeled using four distinct colors. The area is characterized by brief repetitive sequences, and to determine the precise paths for gap-filling, the process of mapping nanopore reads onto this region is utilized. Long reads that are capable of traversing the repetitive structure are then extracted to facilitate path building.

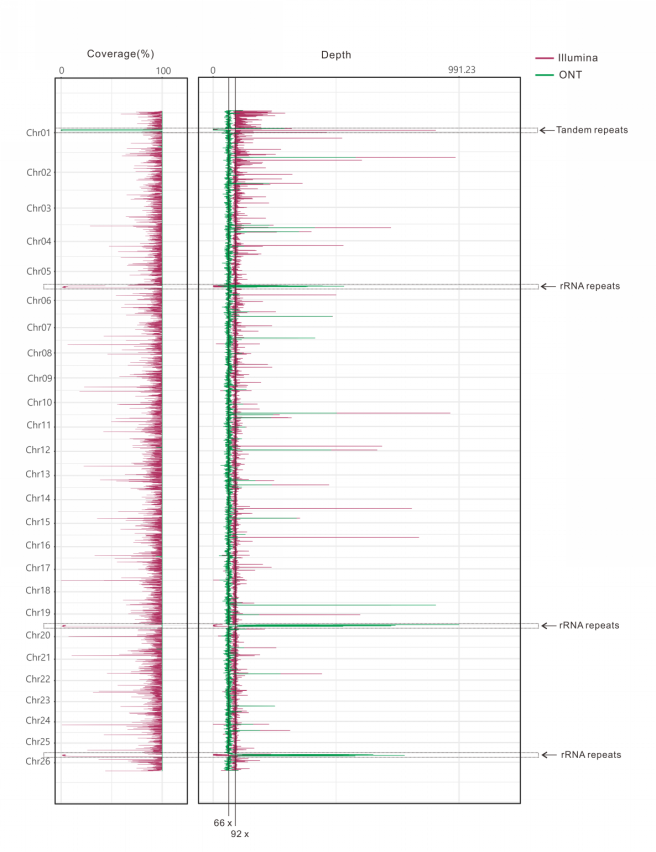

**Supplementary Figure 4. Genome assembly validation achieved by analyzing sequencing coverage and depth in relation to the 26 chromosomes in *P. patens***. The coverage (0-100%) and depth information of Illumina and ONT sequencing reads on 26 chromosomes are illustrated in the left and right images, respectively. The statistical analysis was performed using a nonoverlapping window of 50 kbp. Except for the repetitive region adjacent to the centromere of Chr01, which was deliberately omitted from the secondary mapping findings, the ONT reads demonstrated comprehensive coverage of all other chromosomal regions. Furthermore, the sequencing depth of the multicopy rRNA region was markedly elevated, exceeding the typical chromosome sequencing depth of 66x, which further supports the notion of the presence of several copies of rRNA.

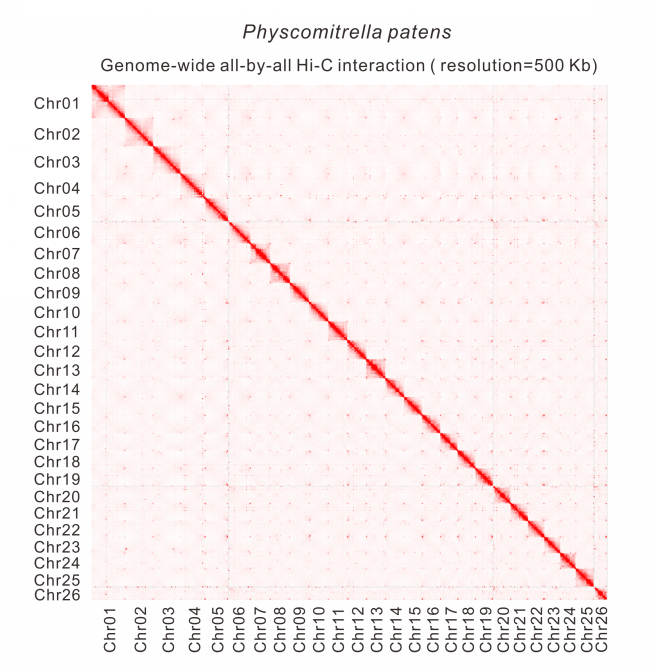

**Supplementary Figure 5. Hi-C interactions among 26 chromosomes at a 500 kbp resolution**.

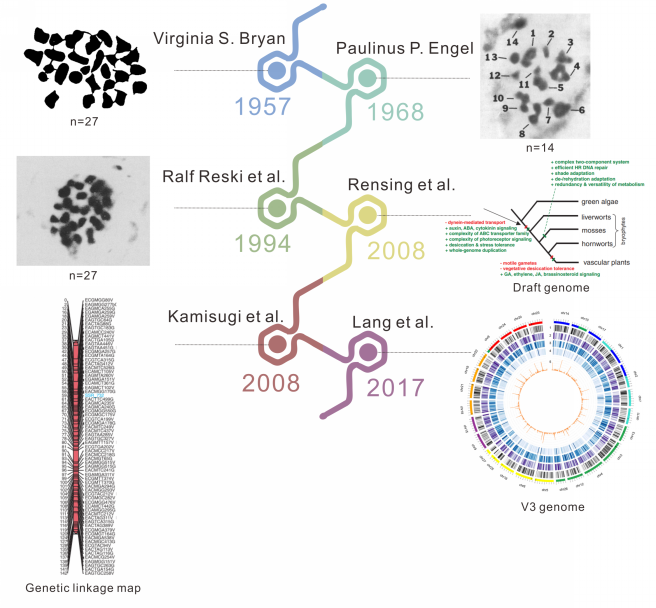

**Supplementary Figure 6. A compilation of pertinent time nodes in the study of chromosome numbers as well as notable research papers that explore the advancements in the field of *P. patens* genome research**.

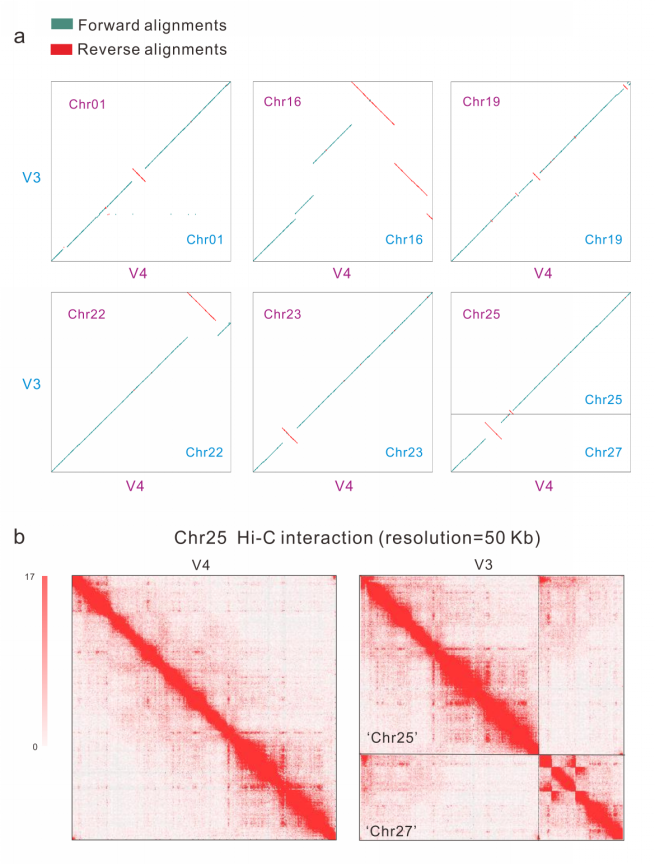

**Supplementary Figure 7**. **Structural conflicts between two distinct versions of particular chromosomes**. **a**, Synteny comparison of six chromosomes between V4 and V3. Malachite green and red dot plots show forward and reverse alignments of the V4 assembly to the V3 assemblies, respectively. **b,** Overview of the Hi-C heatmap for Chr25 between V4 and V3 with a 50 kbp resolution. Notably, two chromosomes (Chr25 and Chr27) in the V3 genome are actually one chromosome corresponding to Chr25 in V4.

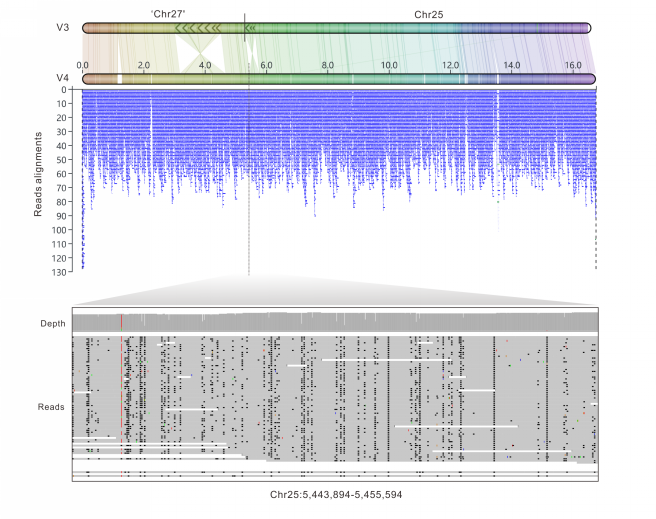

**Supplementary Figure 8. Assembly accuracy validation for Chr25 in V4 by read mapping**. The above depiction aims to compare the level of collinearity displayed by Chr25 in the V3 and V4 versions. The top section of the diagram portrays the amalgamation of two pseudochromosomes in V3, with their boundaries demarcated by a solid black line. The position of the breakpoint in V4 is indicated by a dashed black line. The middle segment of the diagram illustrates the mapping results of nanopore reads (above 10 kbp). The bottom section of the illustration offers a more comprehensive view of the 5 kbp interval encompassing the breakpoint for detailed scrutiny.

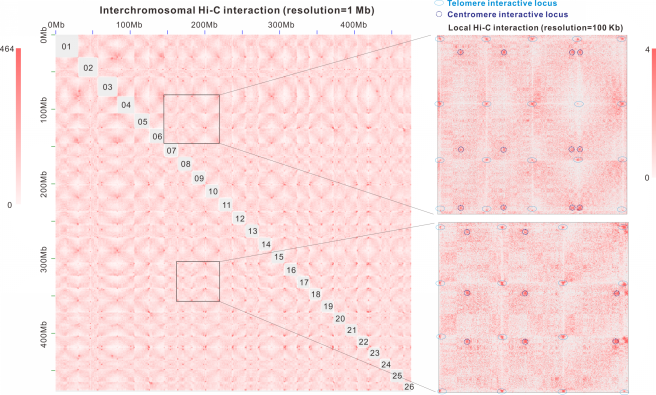

**Supplementary Figure 9. Interchromosomal Hi-C interactions exhibit two types of hotspots according to the positions on each chromosome**. To make these hotspots clearer, intrachromosomal signals were removed. The right panel displays two amplified regions (resolution=100 kbp) that show telomere interactive loci at the borders of chromosomes. Enriched interactions inside chromosomes are presumed to be centromere interactive loci.

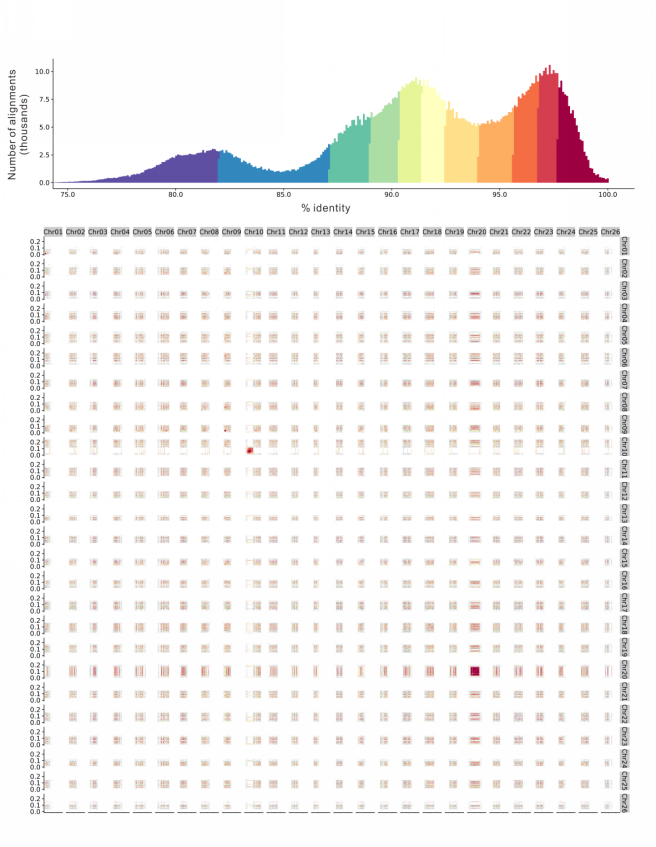

**Supplementary Figure 10. Dot plot for centromere intervals of 26 chromosomes along with their respective upstream and downstream 20 kbp intervals**. The dot plot highlights various arrays of higher-order repeats, which are made evident by the colored percent identity. The similarity-based quantity distribution curve manifests two peaks, each with values greater than 90%. This corresponds with the LTR insertion time distribution expounded upon in the main text.

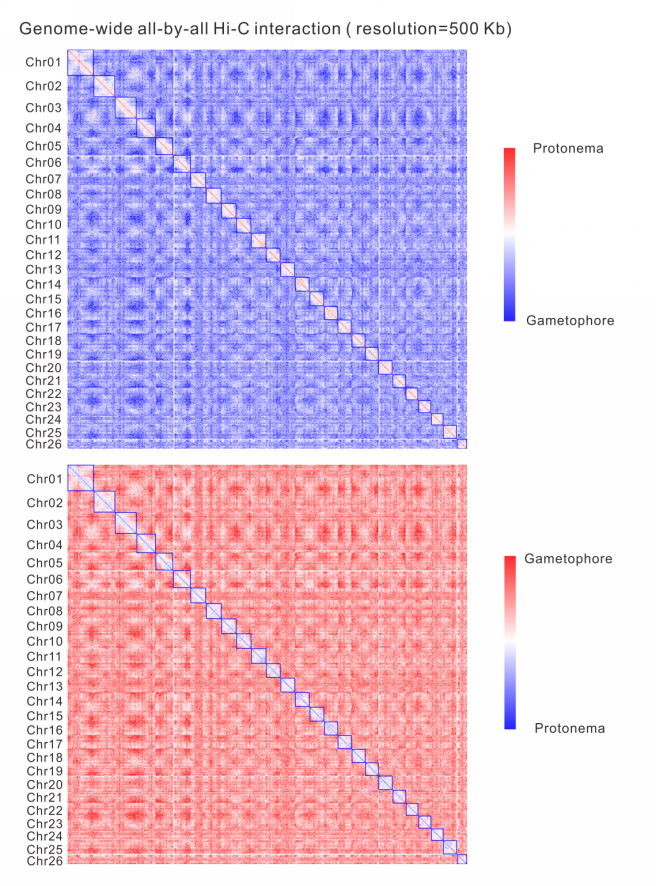

**Supplementary Figure 11. The Hi-C chromatin interaction z score (protonema/gametophore and gametophore/protonema) at a 500 kbp resolution**. The standardized matrix z score heatmap facilitates a comparative evaluation of the interaction intensity between protonema and gametophore chromosomes, both within and across them. Upon detailed examination of the graph, it is evident that the interaction intensity within the protonema chromosomes is significantly higher than that of the gametophore. Conversely, the interaction intensity between chromosomes diminishes.

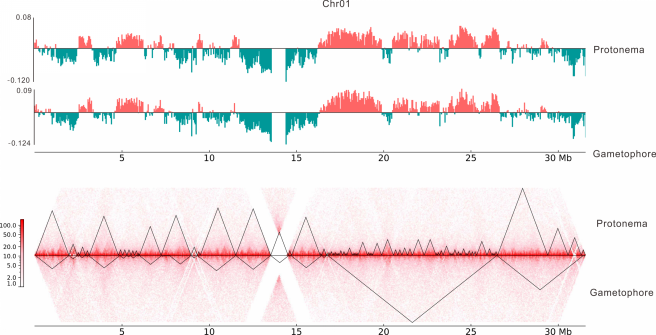

**Supplementary Figure 12. A/B compartment switching and the alteration of topologically associating domain (TAD) boundaries between protonema and gametophore on Chr01**. The bar plots exhibit the fraction of the genome belonging to either the A (depicted in pink) or B (depicted in green) compartment in each of the two tissues that were analyzed. The Hi-C interaction heatmap shows the protonema and gametophore in the upper and lower portions, respectively. The triangular region represents the TAD interval that has been identified.

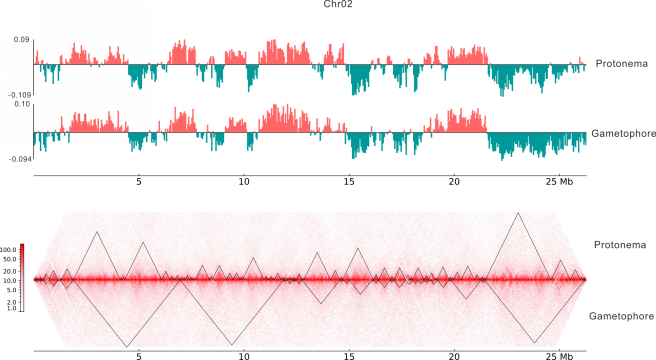

**Supplementary Figure 13. A/B compartment switching and the alteration of topologically associating domain (TAD) boundaries between protonema and gametophore on Chr02**.

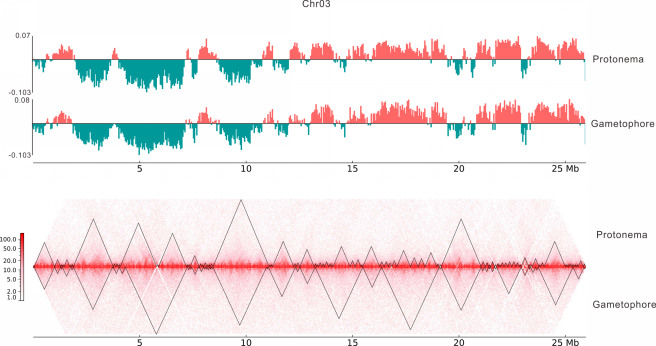

**Supplementary Figure 14. A/B compartment switching and the alteration of topologically associating domain (TAD) boundaries between protonema and gametophore on Chr03**.

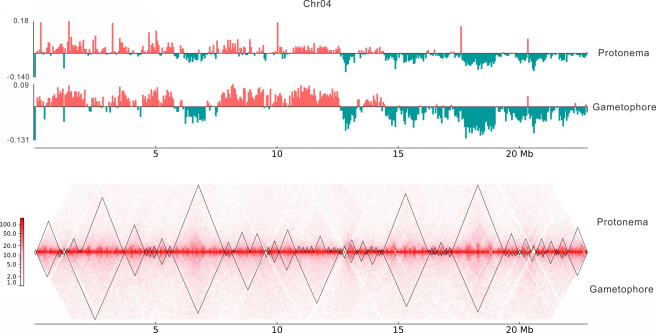

**Supplementary Figure 15. A/B compartment switching and the alteration of topologically associating domain (TAD) boundaries between protonema and gametophore on Chr04**.

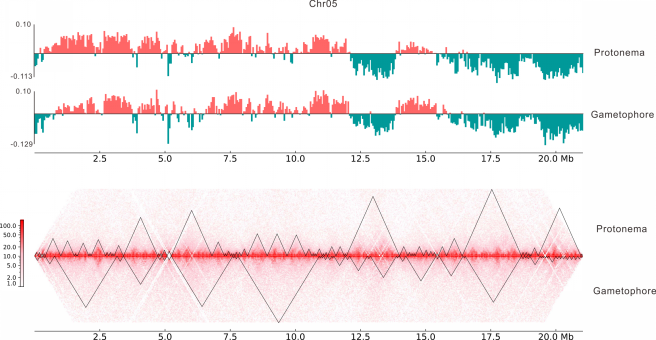

**Supplementary Figure 16. A/B compartment switching and the alteration of topologically associating domain (TAD) boundaries between protonema and gametophore on Chr05**.

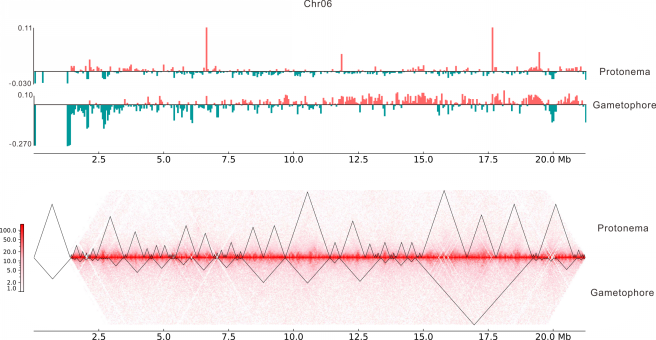

**Supplementary Figure 17. A/B compartment switching and the alteration of topologically associating domain (TAD) boundaries between protonema and gametophore on Chr06**.

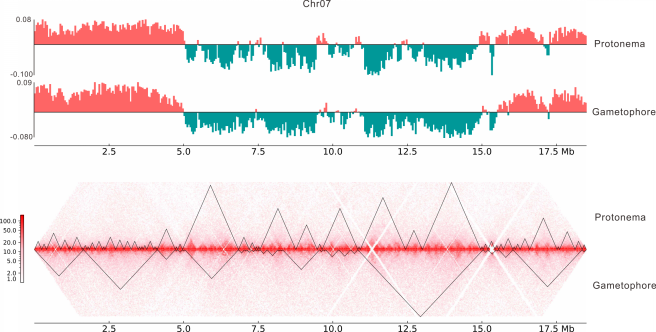

**Supplementary Figure 18. A/B compartment switching and the alteration of topologically associating domain (TAD) boundaries between protonema and gametophore on Chr07**.

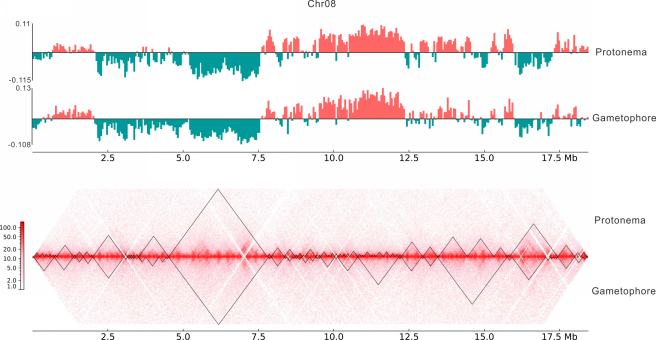

**Supplementary Figure 19. A/B compartment switching and the alteration of topologically associating domain (TAD) boundaries between protonema and gametophore on Chr08**.

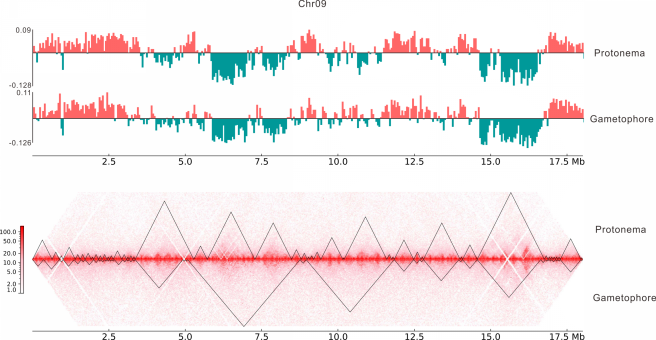

**Supplementary Figure 20. A/B compartment switching and the alteration of topologically associating domain (TAD) boundaries between protonema and gametophore on Chr09**.

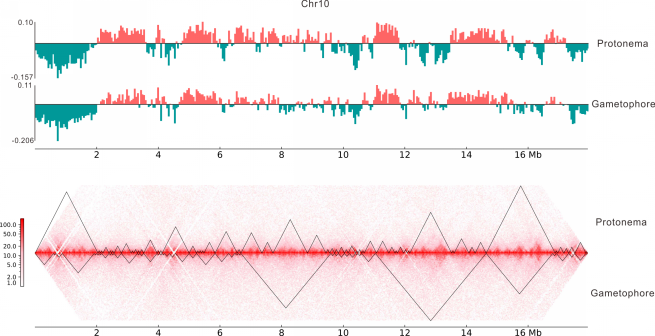

**Supplementary Figure 21. A/B compartment switching and the alteration of topologically associating domain (TAD) boundaries between protonema and gametophore on Chr10**.

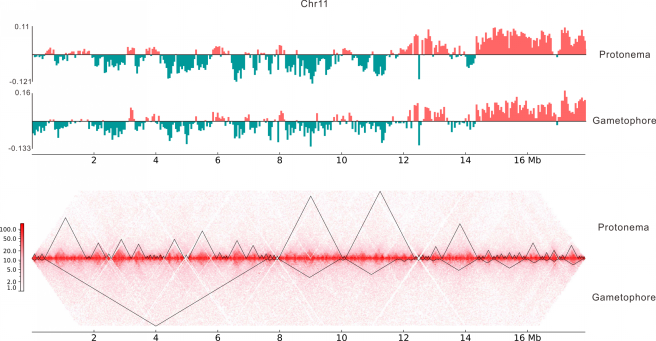

**Supplementary Figure 22. A/B compartment switching and the alteration of topologically associating domain (TAD) boundaries between protonema and gametophore on Chr11**.

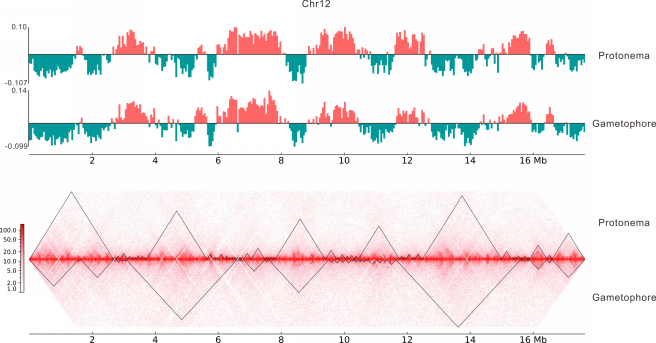

**Supplementary Figure 23. A/B compartment switching and the alteration of topologically associating domain (TAD) boundaries between protonema and gametophore on Chr12**.

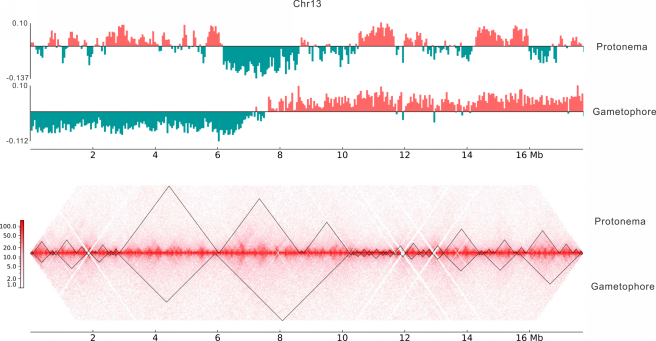

**Supplementary Figure 24. A/B compartment switching and the alteration of topologically associating domain (TAD) boundaries between protonema and gametophore on Chr13**.

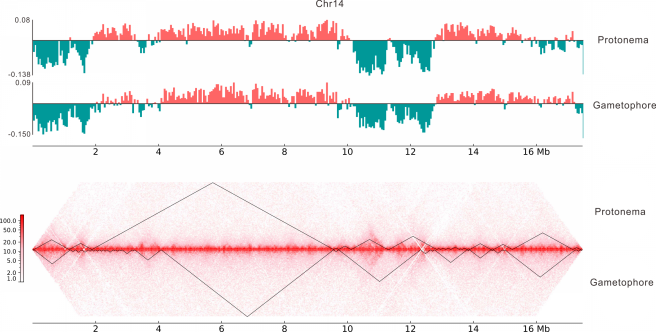

**Supplementary Figure 25. A/B compartment switching and the alteration of topologically associating domain (TAD) boundaries between protonema and gametophore on Chr14**.

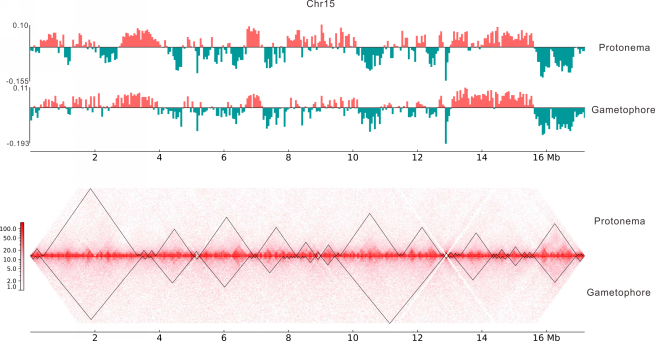

**Supplementary Figure 26. A/B compartment switching and the alteration of topologically associating domain (TAD) boundaries between protonema and gametophore on Chr15**.

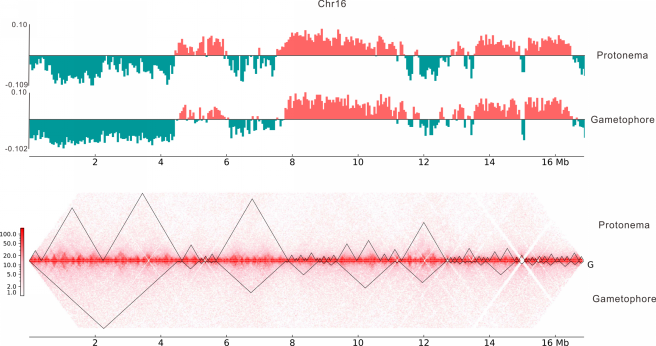

**Supplementary Figure 27. A/B compartment switching and the alteration of topologically associating domain (TAD) boundaries between protonema and gametophore on Chr16**.

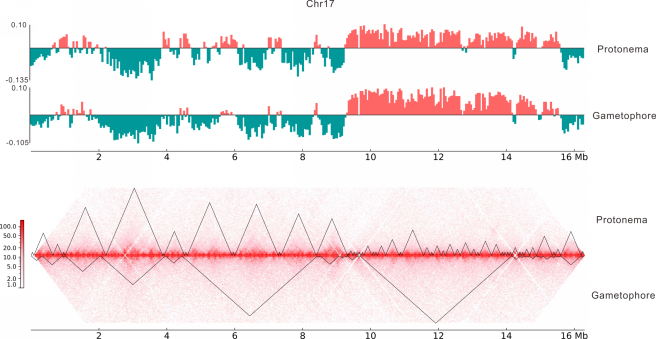

**Supplementary Figure 28. A/B compartment switching and the alteration of topologically associating domain (TAD) boundaries between protonema and gametophore on Chr17**.

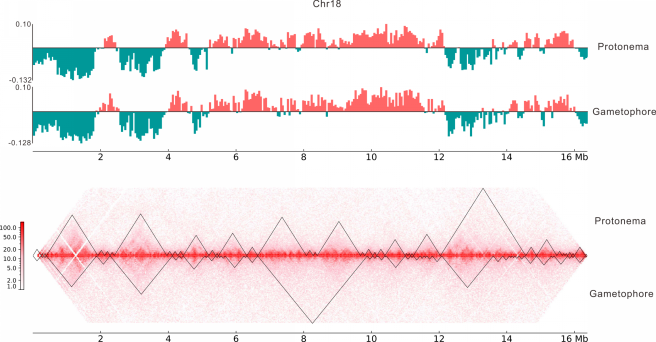

**Supplementary Figure 29. A/B compartment switching and the alteration of topologically associating domain (TAD) boundaries between protonema and gametophore on Chr18**.

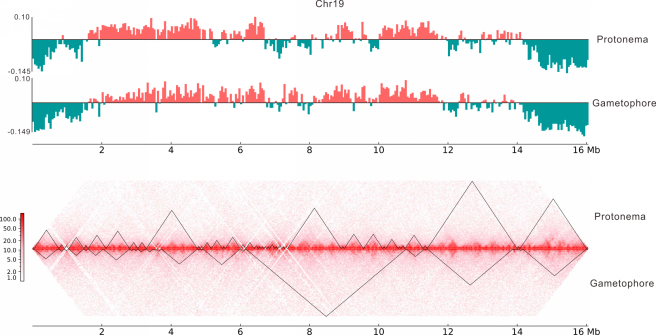

**Supplementary Figure 30. A/B compartment switching and the alteration of topologically associating domain (TAD) boundaries between protonema and gametophore on Chr19**.

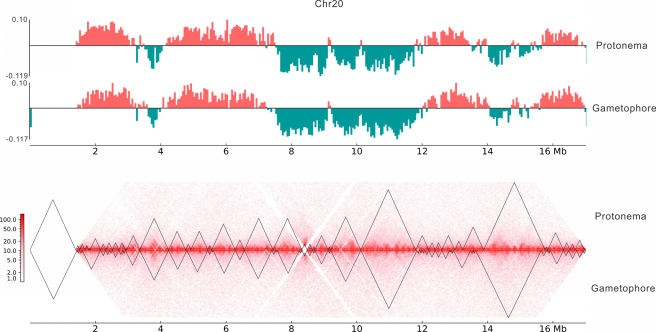

**Supplementary Figure 31. A/B compartment switching and the alteration of topologically associating domain (TAD) boundaries between protonema and gametophore on Chr20**.

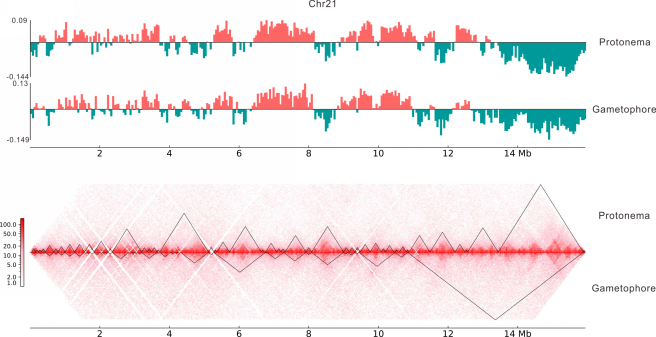

**Supplementary Figure 32. A/B compartment switching and the alteration of topologically associating domain (TAD) boundaries between protonema and gametophore on Chr21**.

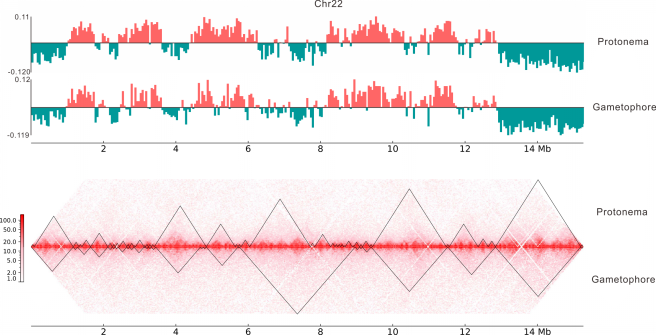

**Supplementary Figure 33. A/B compartment switching and the alteration of topologically associating domain (TAD) boundaries between protonema and gametophore on Chr22**.

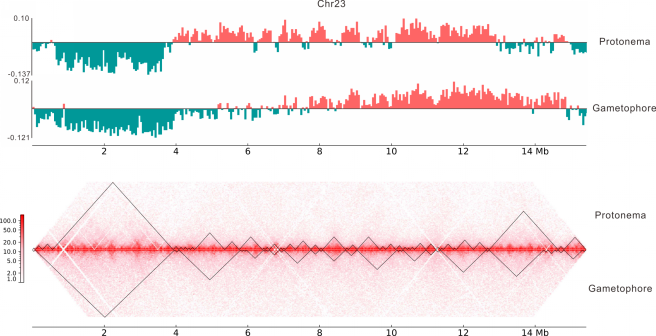

**Supplementary Figure 34. A/B compartment switching and the alteration of topologically associating domain (TAD) boundaries between protonema and gametophore on Chr23**.

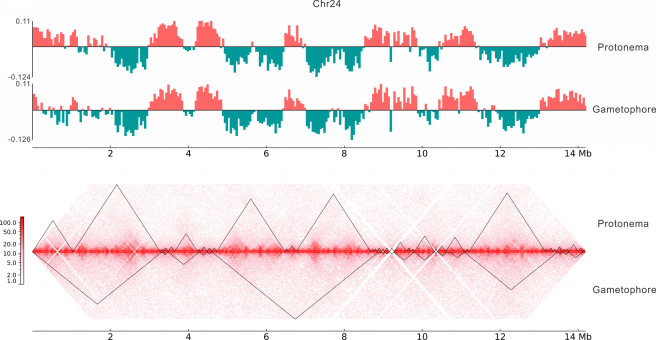

**Supplementary Figure 35. A/B compartment switching and the alteration of topologically associating domain (TAD) boundaries between protonema and gametophore on Chr24**.

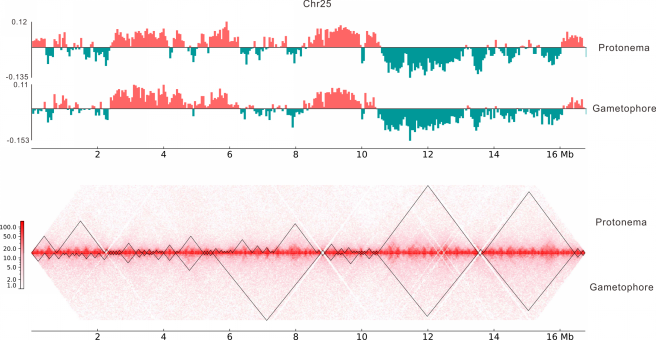

**Supplementary Figure 36. A/B compartment switching and the alteration of topologically associating domain (TAD) boundaries between protonema and gametophore on Chr25**.

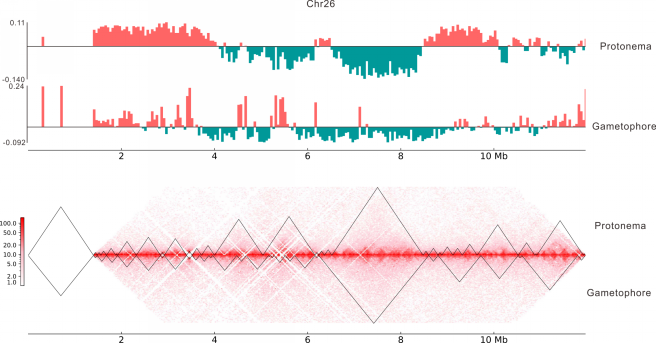

**Supplementary Figure 37. A/B compartment switching and the alteration of topologically associating domain (TAD) boundaries between protonema and gametophore on Chr26**.
